# Moderate intensity aerobic exercise in 6-OHDA-lesioned rats alleviates established motor deficits and reduces neurofilament light and glial fibrillary acidic protein serum levels without increased striatal dopamine or tyrosine hydroxylase protein

**DOI:** 10.1101/2023.07.11.548638

**Authors:** Ella A. Kasanga, Isabel Soto, Ashley Centner, Robert McManus, Marla K. Shifflet, Walter Navarrete, Yoonhee Han, Jerome Lisk, Ken Wheeler, Isha Mhatre-Winters, Jason R. Richardson, Christopher Bishop, Vicki A. Nejtek, Michael F. Salvatore

**Affiliations:** Department of Pharmacology and Neuroscience, University of North Texas Health Science Center, Fort Worth, TX; Department of Psychology, Binghamton University, Binghamton, NY; Department of Environmental Health Sciences, Robert Stempel School of Public Health & Social Work, Florida International University, Miami, FL; Clearcut Ortho Rehab & Diagnostics, Fort Worth, TX

## Abstract

**Background:** Alleviation of motor impairment by aerobic exercise (AE) in Parkinson’s disease (PD) points to a CNS response that could be targeted by therapeutic approaches, but recovery of striatal dopamine (DA) or tyrosine hydroxylase (TH) has been inconsistent in rodent studies.

**Objective:** To increase translation of AE, 3 components were implemented into AE design to determine if recovery of established motor impairment, concomitant with >80% striatal DA and TH loss, was possible. We also evaluated if serum levels of neurofilament light (NfL) and glial fibrillary acidic protein (GFAP), blood-based biomarkers of disease severity in human PD, were affected.

**Methods:** We used a 6-OHDA hemiparkinson rat model featuring progressive nigrostriatal neuron loss over 28 days, with impaired forelimb use 7 days post-lesion, and hypokinesia onset 21 days post-lesion. After establishing forelimb use deficits, moderate intensity AE began 1-3 days later, 3x per week, for 40 min/session. Motor assessments were conducted weekly for 3 wks, followed by determination of striatal DA, TH protein and mRNA, and NfL and GFAP serum levels.

**Results:** Seven days after 6-OHDA lesion, recovery of depolarization-stimulated extracellular DA and DA tissue content was <10%, representing severity of DA loss in human PD, concomitant with 50% reduction in forelimb use. Despite severe DA loss, recovery of forelimb use deficits and alleviation of hypokinesia progression began after 2 weeks of AE and was maintained. Increased NfLand GFAP levels from lesion were reduced by AE. Despite these AE-driven changes, striatal DA tissue and TH protein levels were unaffected.

**Conclusions:** This proof-of-concept study shows AE, using exercise parameters within the capabilities most PD patients, promotes recovery of established motor deficits in a rodent PD model, concomitant with reduced levels of blood-based biomarkers associated with PD severity, without commensurate increase in striatal DA or TH protein.

## Introduction

As ever-increasing evidence supports that aerobic exercise (AE) can slow the rate of progressive motor impairment in Parkinson’s disease (PD) (Ridgel et al., 2009;2019; Schenkman et al., 2018; Shulman et al., 2013; Tihonen et al., 2021; van der Kalk et al., 2019), there is a great need to identify the CNS mechanisms engaged by AE that alleviate motor impairment (Daalen et al., 2022; Li et al., 2023; Salvatore et al., 2022). Identifying these AE-responsive mechanisms is crucial for several reasons. First, a number of barriers to exercise in the PD patient have been identified (Afshari et al., 2016; Horbinski et al., 2021; Schootemeier et al., 2020), including low expectations from practicing exercise, a sedentary lifestyle, lack of social support, depression, fatigue, or the living environment (Ellis et al., 2013; Horbinski et al., 2021; da Silva et al., 2023, Schootemeijer et al., 2020). Moreover, the progressive nature of PD can diminish the physical capabilities needed for beneficial effects of AE (Kanequsuku et al., 2021). Thus, identifying and targeting the AE-responsive mechanisms in CNS by pharmacological or genetic approaches represents a therapeutic strategy to help patients maintain motor function, especially as the disease progresses to advanced stages.

Imaging studies in human PD patients are yielding insight into CNS mechanisms engaged by AE (Fisher et al., 2013; Sacheli et al., 2019; Shih et al., 2019). Rodent PD models represent optimal sources to identify and interrogate which CNS mechanisms could drive motor benefits (Arnold et al., 2016; Castro et al., 2022; Crowley et al., 2019; O’dell et al., 2007; Palasz et al., 2019; Petzinger et al., 2007; 2013; Smith et al., 2011; Tajari et al., 2010; Tillerson et al., 2003). However, while a number of AE-responsive candidate mechanisms have been reported, a single mechanism, or pathway-related cluster of mechanisms, has yet to be revealed as critical for slowing progressive motor impairment. To move forward and fill this knowledge gap, consideration of the conditions of human PD should be imposed in rodent PD models (Barker et al., 2020; Kasanga et al., 2021; Salvatore et al., 2022; Zeiss et al., 2017). First, the physical limitations of the PD patient, a critical and often neglected component in preclinical studies, should be imposed in the AE regimen (Salvatore et al., 2022; Zeiss et al., 2017). Second, AE should be implemented after establishing PD-like motor impairment in the model for 2 reasons; first, PD diagnosis is currently only made after onset of PD cardinal motor signs (Berg et al., 2015, de Lau and Bretler, 2006; Schulman et al., 2011; Tolosa et al., 2021), and second, a substantial percentage of PD patients likely have a sedentary lifestyle (Dontje et al., 2013; Ellingson et al., 2019; von Rosen et al., 2021). Third, longitudinal assessment of AE impact emulates the PD patient practicing AE over months to years, and, in rodent models, reveal the timing of AE impact on motor function to capture mechanistic insight. Fourth, as aging is the number one risk factor for PD, and can limit AE efficacy in parkinsonian rats (Arnold et al., 2017; Kasanga et al., 2021), the neurobiological background of aging represents the age of the typical PD patient (Barata-Antunes et al., 2020; Klasestrup et al., 2022)

We addressed 3 of the 4 elements of AE translation using the established 6-OHDA rat PD model. First, AE frequency (3 times/wk), duration (40 min per session), and intensity (by treadmill speed needed to increase heart rate 60-70% maximum) was applied to be consistent with early-stage PD patients practicing AE at moderate intensity for at least 3 months (Salvatore et al., 2022), which, in both patient and rats, improved motor function (Kasanga et al., 2021). Second, AE was initiated after establishing motor impairment and emulating the severity of tyrosine hydroxylase (TH) protein loss in striatum (>75%) (Kasanga et al, 2022) also occurring in human PD when parkinsonian signs begin (Bernehimer et 1973; Kordower et al., 2013). Third, evaluation of AE impact was done weekly during which time progressive loss of nigrostriatal neurons and hypokinesia appears (Kasanga et al., 2022). To further increase translation of results to human PD (outside of AE implementation criterion), we evaluated serum levels of neurofilament light (NfL) and glial fibrillary acidic protein (GFAP), recently shown to be blood-based biomarkers of disease severity in human PD (Buhmann et al., 2023; Mollenhauer et al., 2020; Tang et al., 2023; Ye et al., 2021; Ygland Rodstrom et al., 2022; Youssef et al., 2023). Our results show motor impairment was alleviated by AE, with gradual recovery beginning after the 2^nd^ week, but without recovery of striatal DA or TH loss. However, AE reduced lesion-elevated serum levels of NfLand GFAP. To our knowledge, this study is the first to report recovery of established motor impairment by an AE regimen that emulates physical capabilities of PD patient, with decreased levels of a peripheral biomarker of PD severity to accompany motor improvement. They also confirm findings from a number of preclinical studies showing motor benefits of AE without increased striatal DA markers.

## Methods

### Animals

Sprague-Dawley rats, purchased from Charles River (Worcester, MA, USA), were used in the exercise study (n=40, male), and in a separate *in vivo* microdialysis study, were used to determine if dopamine (DA) tissue content affected extracellular DA levels at baseline and depolarizing conditions (n=28, 12 male/16 female). Rats were housed under controlled lighting conditions with reverse light-dark cycle so measurements were always recorded in the wake cycle. Standard animal chow and water were available *ad libitum*. All animals were used in compliance with the Office of Laboratory Animal Welfare guidelines and protocols approved by the Institutional Animal Care and Use Committee at the University of North Texas Health Science Center (IACUC-2018-0013), and Binghamton University (IACUC-885-23).

### 6-OHDA lesion induction

Rats were anesthetized with 2-3% continuous inhalation of isoflurane for survival surgery to deliver 6-OHDA. Rats were immobilized in a stereotaxic frame to target the medial forebrain bundle (mfb) at coordinates relative to Bregma (ML -2.0, AP -1.8, DV -8.6) (Paxinos and Watson, 1986). Rats received 6-OHDA (16 μg in 4 μL of vehicle (0.02%w/v ascorbic acid) in the left mfb or vehicle. Vehicle was infused into the contralateral mfb for all 6-OHDA groups and sham groups. The needle was left in place for 10 minutes before removal to allow for maximal toxin diffusion. Body temperature was maintained at 37 °C using a temperature monitor and heating pad.

### Microdialysis

6-OHDA lesioned rats underwent in vivo microdialysis at Day 7 or 28 post-lesion to determine the timing and extent of lesion impact and influence of DA tissue content on extracellular DA levels under basal and depolarizing conditions ipsilateral and contralateral to mfb lesion. The striatum and substantia nigra (SN) were both cannulated during 6-OHDA lesion surgery and targeted either on the side ipsilateral or contralateral to mfb lesion. During the rats’ dark cycle (7:00 A.M. – 1:00 P.M.), microdialysis probes (CMA 12 Elite probe) were inserted into targeted striatal (AP +1.0, ML ± 2.5, DV -3.5) and nigral (AP -5.7, ML ± 2.5, DV -7.0). Rats habituated to the microdialysis chamber for 1 hr prior to a 2-hr baseline collection period. At 170-min into microdialysis, potassium-chloride (KCl) (Tx) spiked aCSF (47.7 mM NaCl, 100 mM KCl, 1.2 mM CaCl2, 1.2 mM MgCl2) was infused into the striatal site for 20 min to evoke DA release. This infusion was replaced with aCSF again for an additional 100 min. Dialysate samples (20 µl) were taken every 20 min for 5 hrs and stored at -80°C until analysis by an ultra-high performance liquid (UHPLC) chromatography. After a 24-hr washout, brains were flash frozen in cold 2-methylbutane (EMD Millipore) for confirmation of striatal placement and dissection of striatal tissue from both hemispheres for subsequent analysis of DA tissue content. Additional information on microdialysis and UHPLC analysis is located in Supplemental Results.

### Confirmation of nigrostriatal lesion phenotype

Forepaw adjustments steps (FAS) was used to measure forelimb use akinesia resulting from DA depletion (Chotibut et al., 2017; Meadows et al., 2017). Briefly, both hind limbs and one forepaw are restrained and the rat is then moved across a table at a speed of 90 cm/10 s, during which the number of adjusting steps taken is counted. 6 trials per forepaw are conducted: 3 forehand trials (lateral steps toward the thumb of the paw that is stepping) and 3 backhand trials (lateral steps towards the pinky of the paw that is stepping), alternating the starting forepaw between rats. The number of steps taken by the forepaw associated with lesioned side is divided by the number of steps taken by the forepaw associated with sham-operated (contralateral to lesion) side and expressed as overall % intact. Our previous work demonstrated that rats with >25% loss of forelimb use with the lesioned side have a statistically significant difference in eventual onset of hypokinesia 21 days after lesion (Kasanga et al., 2022).

### Amphetamine rotation test

Ipsilateral rotations (toward the lesioned side) were counted for 15 min after a 15-min wait period following amphetamine (AMPH) injection (2 mg/kg, i.p.) on day 7 post-surgery. A minimum of 18 rotations (per 15 min) was used as a secondary approach to qualify nigrostriatal lesion.

### Open-field Locomotor Assessment

Open-field locomotor activity chambers (Opto Varimex 4 Animal Activity Monitoring System, Columbus Instruments, Columbus, OH, USA) were used to quantify locomotor activity as total distance traveled (DT), as total cm in one hr, at the following 6 designated time points; pre-acclimation to exercise, post-acclimation to exercise, and 7, 15, 22, and 29 days post-lesion.

### Apomorphine rotation

Apomorphine (0.05 mg.kg, s.c.) was administered immediately after the 1 hr open-field locomotor activity assessment on the final day of the experiment. Rotations contralateral to lesion were counted for the first 30 min post-injection by an observer blind to experimental group.

### Exercise regimen

There were two main groups-a control group that underwent sham surgery and a 6-OHDA group. Within each group, there was an exercise (EX) and non-exercise (NE) group. Power analysis using historical data (effect size = 1.126, alpha = 0.05, Power = 0.60) indicated a sample size of 8. To increase power and account for attrition due to surgery-related issues, 6-OHDA groups had a sample size of 12.

All rats underwent three main phases in the regimen (Fig. 1). The first phase was treadmill acclimation which was preceded by establishment of baseline locomotor testing from the FAS and open-field locomotor tests. Acclimation consisted of introducing the rats to increasing speeds of the treadmill (without footshock) and ensured all rats experienced the treadmill environment (Arnold and Salvatore, 2014). A post-exercise acclimation assessment of locomotor testing was completed. This was followed by 6-OHDA lesioning or sham surgery. Seven days after 6-OHDA surgery, lesion efficacy was verified using the FAS and amphetamine rotation test. Rats were randomly subdivided into the NE or EX groups, notably ensuring that assignment of lesioned rats (exceeding more than 25% loss of forelimb use in lesioned forelimb vs intact forelimb) showed comparable overall lesion efficacy between the lesion groups, and compliance to exercise (Arnold and Salvatore, 2014).

**Figure 1.**
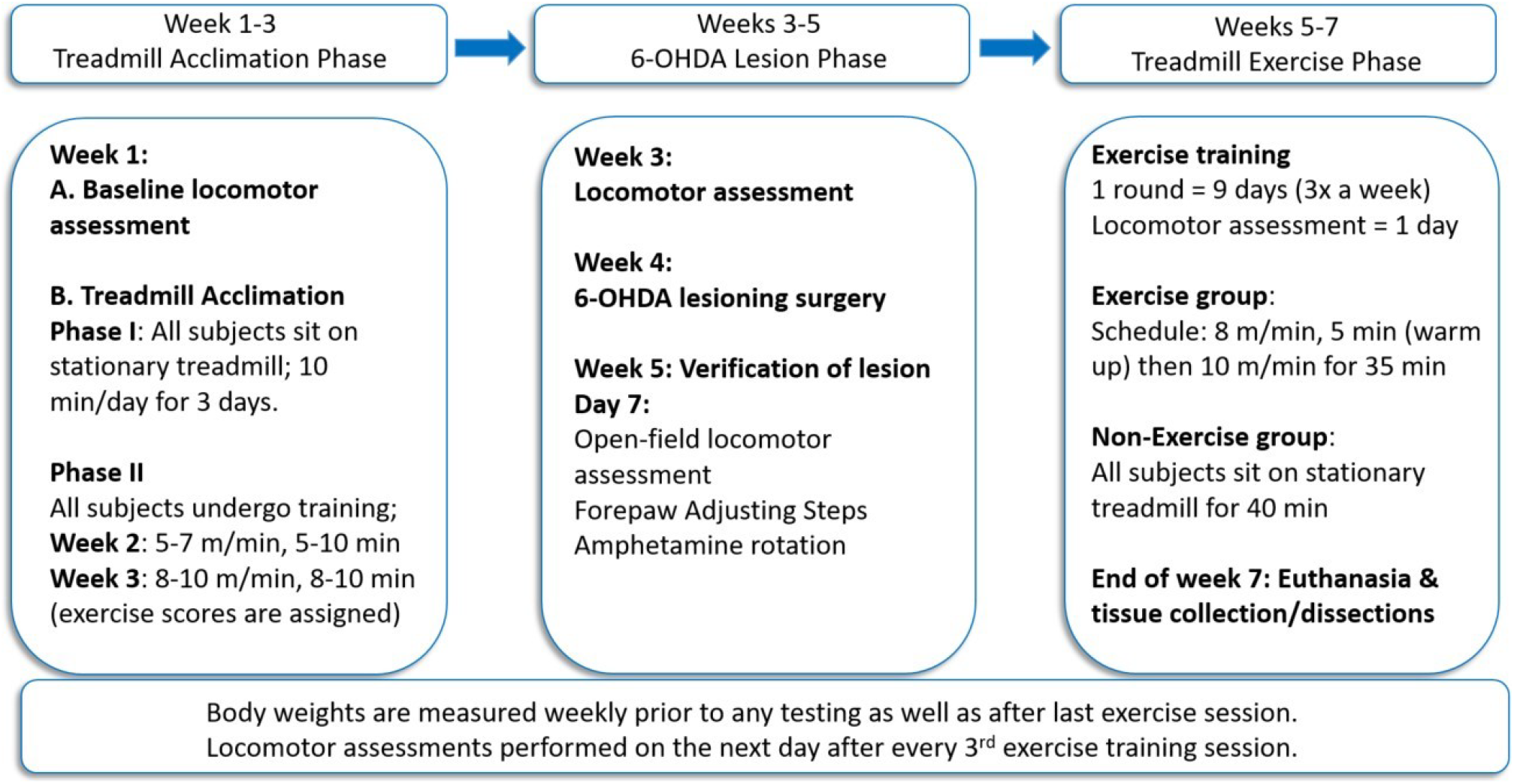
Experimental timeline. Locomotor testing was conducted on all rats at 2 different time points, before and after treadmill exposure and training, before 6-OHDA lesion induction or sham-operation. 7 days after lesion, forepaw adjusting steps (FAS) qualified lesion efficacy (min 25% loss of forelimb use), and rats were randomly placed into non-exercise or exercise groups ensuring no significant differences in FAS between groups. Treadmill exercise was conducted 3 times a week for 3 weeks, with locomotor assessments conducted one day following the last of the 3 exercise sessions each week. On the day of the final locomotor assessment, which included the apomorphine rotation test, rats were euthanized for tissue dissection.

Exercise began on days 8-10 post-lesion (or sham surgery), 1-3 days after establishing decreased forelimb use, and continued for 3 weeks, consisting of 9 total exercise sessions, 3 times a week, 40 min on alternate days. This frequency was chosen as it has been shown by Pelosin et al (2017) and others (Li et al., 2023; Osborne et al., 2022; Salvatore et al., 2022; Silveira et al., 2018; van der Kolk et al., 2019) that in PD patients, this is a commonly used frequency each week for patients, and in some cases, portend improvement in Unified Parkinson’s Disease Rating Scale scores or quality of life. The non-exercise group was placed on a stationary treadmill for the duration of the exercise session. Body weight was tracked weekly and locomotor performance was assessed the day after every 3rd exercise training session. Rats were euthanized on the same day as their 3^rd^, and final, post-exercise locomotor assessment immediately after apomorphine rotation test.

### Body weight measures

Weight of the test subjects was taken to evaluate effects of the AE regimen and determine any possible relationship or influence upon locomotor activity measures.

### Tissue dissection, processing, and DA and DA turnover analysis

Rats were euthanized after the last locomotor session by decapitation following brief exposure to isoflurane. The striatum was dissected on wet-ice and immediately cooled on dry-ice and saved for subsequent ultra high-performance liquid chromatography (UHPLC) analysis of DA tissue content, DA turnover, and RT-qPCR assay of tyrosine hydroxylase (TH) mRNA and quantitative western blot of TH protein (as ng TH / µg total protein) (Salvatore et al., 2012).

In both studies, fresh frozen tissue was sonicated in ice-cold 0.1M perchloric acid solution and centrifuged. Supernatants were aliquoted for analysis of DA and DOPAC (10 μL per injection volume). The resulting protein pellet was further sonicated in 1% sodium dodecyl sulfate solution (with 5 mM Tris buffer, pH 8.3 & 1 mM EDTA) to determine total protein using the bicinchoninic acid method, to which DA tissue content was normalized after analysis of DA peak height by UHPLC (ThermoFisher Scientific). Additional details for UHPLC analysis are reported in Supplemental Results.

### mRNA extraction and RT-qPCR

RNA was extracted from tissue dissections to evaluate TH mRNA using RNeasy® Micro kit (QIAGEN). The tissue was homogenized, followed by cDNA synthesis reaction performed with iScript™ cDNA Synthesis Kit using the resulting RNA. A cDNA optimization was performed on pooled cDNA samples in differing dilutions and side (lesioned and contralateral to lesion) to determine the optimal load for the RT-qPCR reaction.. The primer mixes used for this experiment were the PrimePCR™ SYBR® Green Assay: TH, Rat for the gene of interest and the PrimePCR™ SYBR® Green Assay: GAPDH Rat for the control gene. Samples were run in duplicate using each primer mixture for comparison. Additional details for cDNA synthesis and PCR cycling program used are reported in Supplemental Results.

### Determination of total protein expression in CNS tissues

SDS gel electrophoresis was conducted after total protein assay (Salvatore et al., 2012).

After protein transfer, nitrocellulose membranes were stained with ponceau S and imaged for normalization of total protein, blocked for > 2 hours, and placed in respective primary antibody for 2 hours. The specific antibody used for TH was purchased from Millipore ((cat #: AB152; 1:1000 dil), Temecula, CA). The membranes were then exposed to secondary antibody and imaged using BioRad imager V3 Chemi-Doc Touch.

The Image Lab software (Bio-Rad Life Sciences, CA) was used for analysis of protein expression. For total TH assessment, the amount of TH, in nanograms (ng) per microgram (μg) protein, was interpolated from standard curves as previously described (Salvatore et al., 2012; Salvatore and Pruett, 2012; Kasanga et al., 2019).

### Serum analysis of biomarkers

Neurofilament light chain (NfL) and glial fibrillary acidic protein (GFAP) levels were quantified in the serum of rats, processed from blood collected at tissue dissection, using the R-PLEX Human Neurofilament Light Assay (cat# K1517XR-2; Meso Scale Discovery, Rockville, MD, United States) and the R-PLEX Human GFAP Assay (cat# K1511MR-2; Meso Scale Discovery, Rockville, MD, United States), respectively. Assays were validated for sample compatibility and accurate analyte measurement. Serum samples from all experimental groups were run in duplicates and incubated overnight. All other directions were followed according to the manufacturer’s protocol. Plates were read on MESO QuickPlex SQ 120 and analyzed using the DISCOVERY WORKBENCH analysis software (version 4.0).

### Statistics

Exclusion criterion flow-chart for statistical assessments of the motor and neurochemistry outcomes are presented in Suppl. Table 1. With the aforementioned forelimb deficit exclusion criterion, 1 rat each from the NE (1/10) and EX (1/11) groups did not meet the >25% loss of forelimb use, but had >90% striatal DA and TH protein loss. All other rats with >25% loss of forelimb use prior to exercise (NE; *n=* 9/10; EX, *n=* 10/11) also had >90% striatal DA or TH protein loss. There were 2 rats in the NE and 1 rat in the EX group that did not meet the 25% forelimb loss threshold and all 3 rats averaged 25% DA loss.

Evaluation of AE impact on locomotor functions used repeated measures 3-way and 2-way ANOVA, matching raw value or calculated results obtained at each time point (post-exercise acclimation (PA) day 7, 15, 22, and 29 post-treatment ((Tx) lesion or sham-operation) for use of forelimb associated with lesion versus use of intact limb and open-field locomotor activity. Calculated results used the raw results obtained at post-acclimation to exercise to determine impact of Tx over weeks post-Tx, or the raw results obtained after completing the first week of exercise to control for any possible exercise-related fatigue. In both calculations, these values served as the denominator to determine % change during lesion progression or sham-operation effects with and without exercise. Apomorphine rotations were evaluated with the non-parametric Mann-Whitney test, as results were not normally distributed in the lesioned groups.

In the 3-way ANOVA, the independent variables were 1) number of weeks post treatment (Tx) (sham-operation or lesion), 2) sham or lesion, and 3) NE or EX. After determining if there was a difference between sham and lesioned groups (including interactions with exercise or weeks post-Tx), the sham and lesioned groups were analyzed separately using repeated measures 2-way ANOVA. With evidence of a significant experimental effect (EX vs NE) or an interaction with assessment period, post-hoc tests were Bonferroni’s multiple comparison test or t-test, were used as indicated. Evaluation of extracellular DA at baseline (BL) or depolarization-stimulated DA by KCl used 3-way ANOVA, with independent variables being 1) weeks post-lesion, 2) contralateral vs ipsilateral side dialysate collection, and 3) BL vs KCl, after meeting criterion of ≥80% DA tissue content loss by lesion. This was followed by 2-way ANOVA analyzing BL and KCl-obtained results separately.

Evaluation of AE impact on DA tissue content and TH mRNA and TH protein used repeated measures 2-way ANOVA, matching lesioned or sham-operated sides against respective contralateral (sham-operated or intact) sides, respectively. Biomarker analysis of GFAP and NfL was done using 2-way ANOVA, with independent variables being lesion v sham, and exercise v non-exercise., followed by unpaired t-test for post-hoc comparisons. The Grubb’s test identified outliers with 95% confidence, using alpha=0.05 for the respective n for each dependent measure for each group.

## Results

### Impact of 6-OHDA lesion on extracellular DA

Loss of TH neurons is progressive between 7 and 28 days with this 6-OHDA regimen, but loss of striatal DA tissue content and TH protein are exceed 90% by day 7, with decreased forelimb use (Kasanga et al., 2022). Analysis of extracellular DA by 3-way ANOVA revealed highly significant differences between ipsilateral and contralateral to lesioned sides (F_1,12)_ =76.7, *p* <0.0001) and BL vs KCl conditions (F_1,12)_ =37.4, *p* <0.0001). This 6-OHDA regimen (Fig. 2 A,B), substantially reduced extracellular DA levels by >90% under depolarizing conditions (KCl), but not baseline (BL) conditions (Lesion x BL vs KCl, (F_(1,12)_ =35.3, *p*<0.0001), ipsilateral to lesion at day 7 and 28 (Fig. 2B). There was no influence of time after lesion (F_(1,12)_ =0.001, *p*=0.99) under baseline (Fig. 2A) (although highly variable at both time points contralateral to lesion) or depolarizing conditions contralateral to lesion ((F_(1,12)_ =0.001, *p*=0.99) (Fig. 2B). DA tissue content comparison against the extracellular DA levels was >90% loss at both time points, indicating >90% loss of striatal DA tissue content greatly reduces DA release capacity by the same magnitude. These results confirm DA signaling in striatum is severely compromised one day prior to AE intervention.

**Figure 2.**
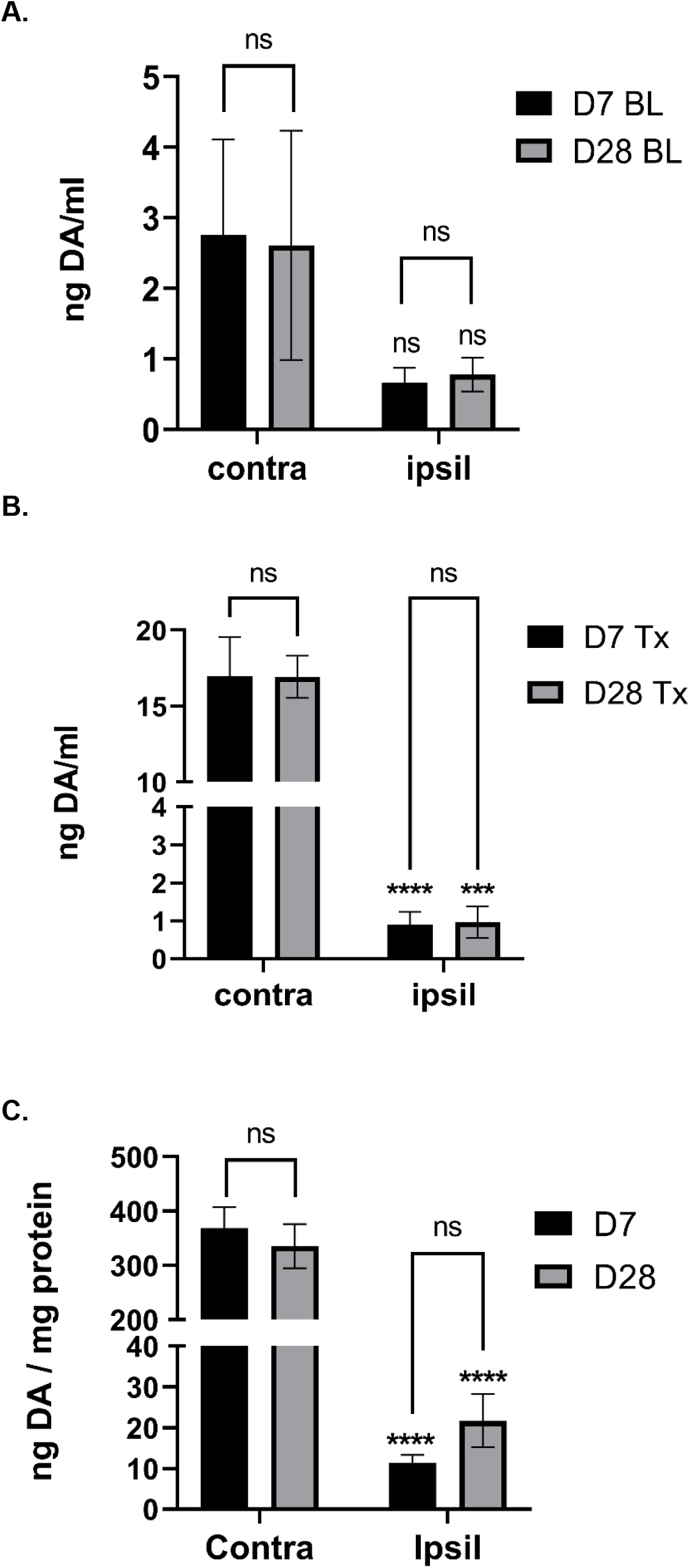
Extracellular DA release capacity is tied to DA tissue content. Extracellular DA under baseline (BL) and depolarizing conditions (Tx) during lesion progression. **A.** Extracellular DA under BL conditions is not significantly affected by lesion (F_(1,12)_ = 2.59, *p=*0.13), or influenced by time past lesion (F_(1,12)_ = 0.01, *p=*0.98). **B.** Extracellular DA under Tx is significantly affected by lesion (F_(1,12)_ = 74.9, *p*<0.0001), but not influenced by time past lesion (F_(1,12)_ = 0.001, *p=*0.98). Contra vs. Ipsil; Day 7 (t=6.58, \*\*\*\**p*<0.0001, df=12), Day 28 (t=5.74, \*\*\**p*=0.0002, df=12), **C. DA tissue content.** DA tissue content is significantly affected by lesion (F_(1,13)_ = 163, *p*<0.0001), but not influenced by time past lesion (F_(1,13)_ = 0.67, *p=*0.43). Contra vs. Ipsil; Day 7 (t=10.1, \*\*\*\**p*<0.0001, df=13), Day 28 (t=8.1, \*\*\*\**p*<0.0001, df=13).

### Locomotor function during exercise acclimation

All rats acclimated to the treadmill environment and speed of the AE regimen prior to random assignment into experimental groups. Although AE acclimation affected locomotor activity (F_(1,36)_=11.8, *p* =0.0015) overall, there was no significant difference between rats subsequently assigned into sham and 6-OHDA lesion groups ((F_(1,36)_=0.001, *p* =0.97) or between 6-OHDA lesioned group rats randomly designated for the NE or EX groups (F_(1,22)_=1.66 *p* =0.21). AE acclimation had negligible effect on forelimb use (Suppl Fig. 1).

### Locomotor function during nigrostriatal lesion progression

Prior to assignment into NE or EX groups, reduced forelimb use was confirmed on day 7(1 wk) post-lesion (F_(1,29)_=64.6, *p* <0.0001), with highly significant difference between sham and lesion groups (Sham, 105.7 ± 2.6%; 6-OHDA, 46 ± 4.3%) (F_(1,31)_=97.8, *p* <0.0001), but not between rats assigned to NE or EX groups prior to initiating exercise (F_(1,31)_=0.13, *p*=0.72) (Fig. 3A). There was also no significant difference in AMPH rotations between the rats assigned to NE and EX groups (Suppl. Fig 2).

**Figure 3.**
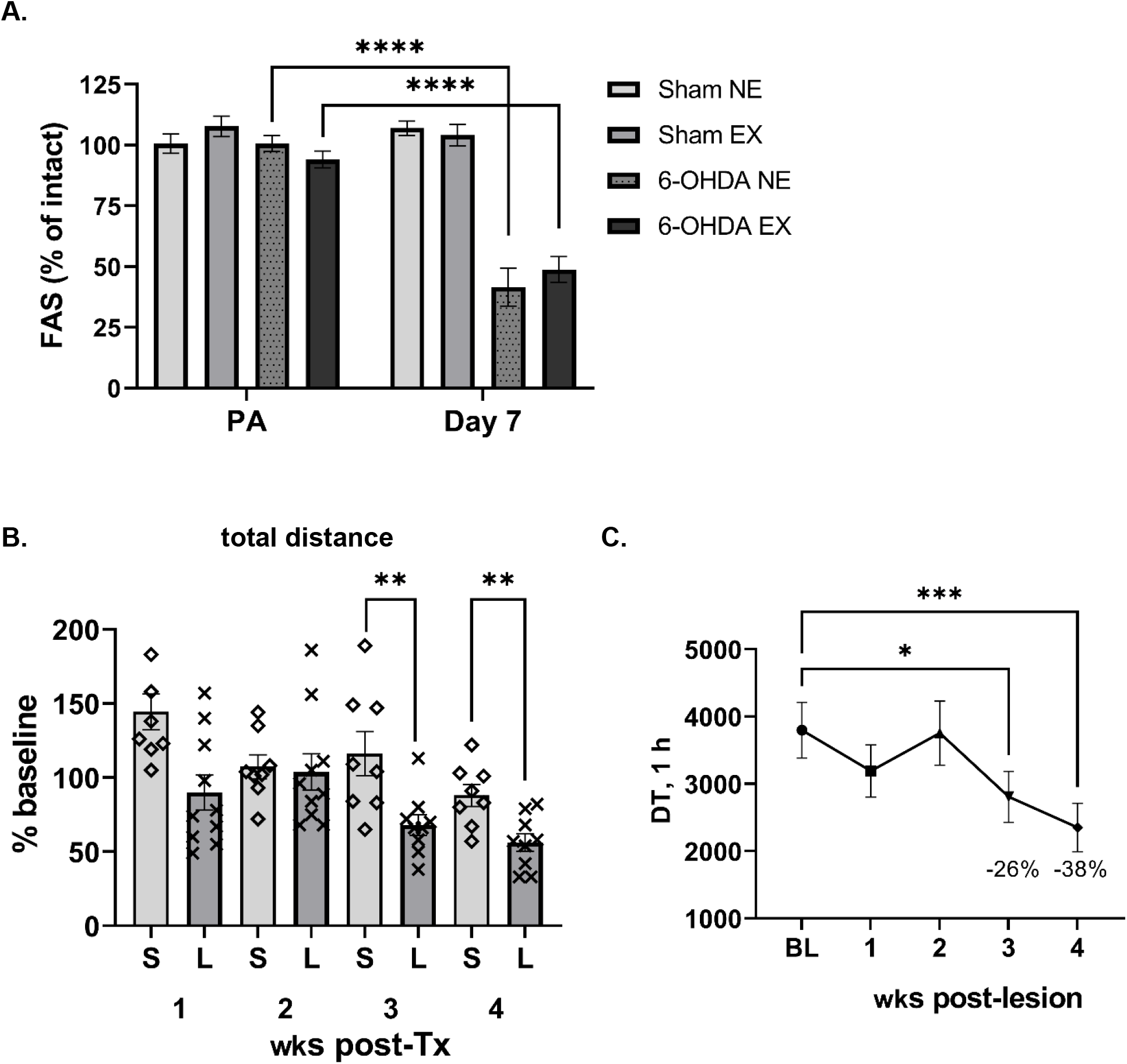
Timing of 6-OHDA lesion impact on locomotor function. A. Establishment of forelimb use deficits. Use of the forelimb associated with the 6-OHDA lesion, but not sham-operation, decreased ∼50% 7 days after lesion induction (Lesion x post treatment, F_(1, 29)_ = 71.91, *p* < 0.0001)); Post-acclimation (PA) v Day 7 (Sham NE, t=0.99, ns; Sham EX t=0.52, ns; 6-OHDA NE, t=9.12, \*\*\*\**p*<0.0001; 6-OHDA EX, t=8.28, \*\*\*\**p*<0.0001). **B. Timing of hypokinesia onset from 6-OHDA lesion against sham-operation.** 6-OHDA lesion significantly decreased locomotor activity (as total distance/hr) in the NE group beginning 3 wks post-lesion. Lesion v sham x weeks post-Tx (F_(3, 46)_ = 2.90, *p* = 0.045); Sham (S) v 6-OHDA (L) (mean ± SEM), 1 wk (144 ± 12% v 90 ± 12%; t=3.17, \*\**p*=0.006, df=16); 2 wk (107 ± 8% v 90 ± 12%; t=0.23, *p*=0.82, df=16); 3 wk (116 ± 15% v 68 ± 7%; t=3.05, \*\**p*=0.008, df=15); 4 wk (88 ± 7% v 56 ± 6%; t=3.37, \*\**p*=0.004, df=15); **C. Hypokinesia onset in 6-OHDA lesion group.** In the NE group, lesion reduced locomotor activity (as distance travelled (DT) in 1 hr) at 3ks post-lesion. One way ANOVA, F_(4,36)_=6.91, *p*=0.003. BL v wk 1 (q=1.82, *p*=0.23); BL v wk 2 (q=0.12, *p*=0.99); BL v wk 3 (q=2.97, \**p*=0.018); BL v wk 4 (q=4.34, \*\*\**p*=0.0004). Dunnett’s multiple comparisons test.

Hypokinesia was confirmed at 3 wks post-lesion, with a significant decrease relative to post-acclimation locomotor activity as compared with sham-op group (F_(1,16)_=11.2, *p* =0.004), weeks post-treatment (F_(3,46)_=7.84, *p* =0.0003) (Fig 3B). Distance covered in the open-field was significantly reduced in the lesioned NE group by 3 wks (Fig. 3C).

Sham-operation produced a transient increase in locomotor activity respective to baseline and greater than changes relative to baseline in the lesion group 1 wk after lesion (Fig. 3B).

### Recovery of forelimb use after exercise

After confirming comparable reduction in forelimb use 1 wk after 6-OHDA lesion, (and no difference in AMPH rotations (Suppl Fig 2)), rats placed into the EX group showed gradual recovery of forelimb use on the lesioned side at the end of the 2^nd^ wk (exercise, F_(1,16)_= 5.49, *p*=0.032) that was maintained as weeks of post-lesion increased (F_(3,47)_= 4.14, *p*=0.011) (Fig. 4A)). Exercise did not affect forelimb use on the intact side (Suppl Fig 3), indicating that AE-related recovery was due to increased use on the lesioned side.

**Figure 4.**
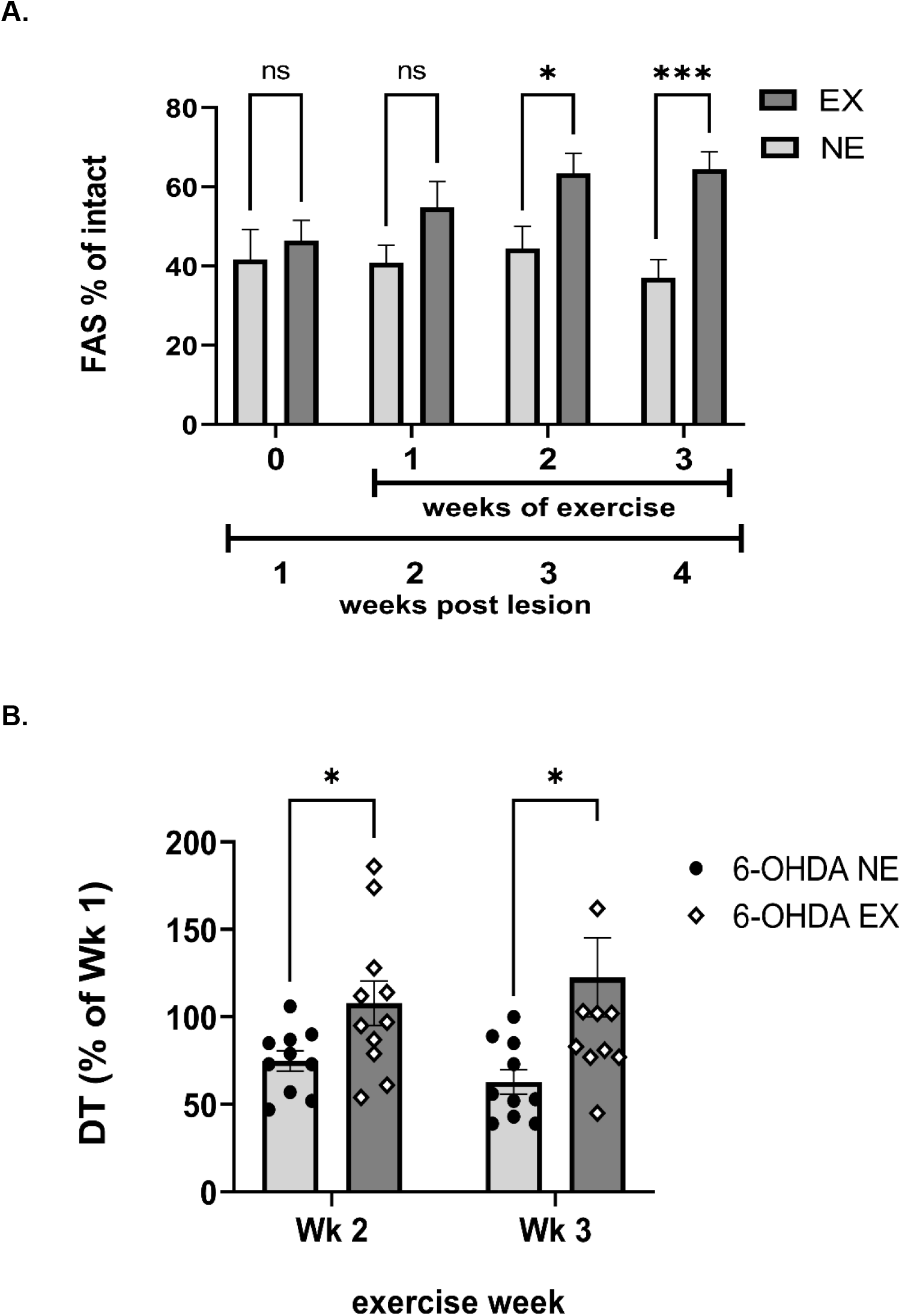
Exercise recovery and prevention of locomotor impairment. A. Recovery of 6-OHDA lesion established forelimb use deficit. Deficit in forelimb adjustment steps (FAS) taken on lesioned side averaged ∼50% of FAS taken on the intact side 1 wk after lesion, prior to initiation of exercise. Exercise promoted recovery of forelimb use, with significant differences by the end of the 2^nd^ week of exercise. Weeks post-lesion x exercise (F_(3, 47)_ = 5.52, *p*=0.003). Post-hoc unpaired t-test results (as NE v EX, % of intact, t-ratio, *p*-value) for each week of exercise. Week 0 (day 7 post lesion), (assigned to NE, 41.6% v assigned to EX, 46.4%, t=0.54, *p*=0.60, df=15); Week 1 (NE, 40.8% v EX, 54.8%; t=1.66, *p*=0.116, df=16); Week 2 (NE, 44.5% v EX, 63.5%; t=2.57, \**p*=0.021, df=16); Week 3 (NE, 37.0% v EX, 64.4%; t=4.45, \*\*\**p*<0.001, df=16). **B. AE prevents hypokinesia onset.** Locomotor activity did not decrease in the EX group during the exercise period following lesion. Exercise (F_(1, 19)_ = 6.00, *p*=0.024). Wk 2 (t=2.29, \**p*=0.034, df=19); Wk 3 (t=2.43, \**p*=0.026, df=18).

### Prevention of hypokinesia onset during exercise

Locomotor activity changed during the course of the study (weeks of study, F_(4,132)_=17.3, *p* < 0.0001) and there was a highly significant influence of AE (weeks of study x exercise, F_(4,132)_=5.19, *p* = 0.0006). During the study, there was a significant effect of 6-OHDA lesion on locomotor activity (lesion v sham, F_(1,33)_=7.14, *p* =0.012) with a highly significant influence of study duration (weeks of study x lesion v sham, F_(4,132)_=7.21, *p* < 0.0001).

Although there was no influence of exercise on locomotor activity in the sham-op group (exercise, F_(1,14)_=0.66, *p* =0.42; exercise x weeks post-sham-op, F_(3,42)_=0.47, *p* =0.70), there was evidence of exercise-related fatigue after the first week of exercise. Locomotor activity decreased the day after conclusion of exercise in sham (Suppl Fig 4A) and 6-OHDA-lesioned groups (Suppl Fig 4B). Given that the NE group in the 6-OHDA-lesioned cohort did not show any decline until day 21, the possibility of fatigue influence was addressed by evaluating locomotor activity in the final 2 wks of exercise against activity levels after the first week of exercise.

Reduced forelimb use ≥ 25% 1 wk after lesion portends eventual hypokinesia 2 weeks later (Kasanga et al., 2022). During the final 2 wks of the study, there was a significant decrease in locomotor activity in the NE group (F_(2,38)_=7.82, *p* =0.002) (Fig. 3C). During this time, there was a highly significant influence of AE (wks of study x exercise F_(2,38)_=6.82, *p* =0.003). Thus, while there was a continuous decrease in locomotor activity in the NE group during this time, no decrease occurred in the EX group (exercise; F_(1,19)_=6.00, *p* =0.024) (Fig. 4B).

### Apomorphine rotation

6-OHDA lesion increased apomorphine rotations compared to sham-operation groups (F_(4,30)_=19.6, *p=*0.0002), with significantly less contralateral rotations (mean ± SEM) in the lesioned EX group (65 ± 24) compared to the lesioned NE group (175 ± 31) (Suppl Fig 5).

*Body weight and exercise*.

6-OHDA lesion affected body weight without any difference between the NE and EX groups. AE in the sham-operation group increased weight (Suppl Fig. 6). There was no significant relationship between baseline body weight or any of the 3 locomotor parameters (FAS, distance, speed) before or after exercise (Suppl Table 2), indicating that recovery or preservation of motor function was not associated with changes in body weight.

### Serum levels of NfL and GFAP

Analysis of blood collected during tissue dissection showed two highly significant effects on serum levels of Nfl (Fig. 5A) and GFAP (Fig. 5B). First, with verified DA and TH protein loss of ≥75% after 6-OHDA lesion, NfL serum levels increased 40% (mean ± SEM) sham NE = 232 ± 11, 6-OHDA NE = 328 ± 20 (F_(1,29)_ = 43.3, *p*<0.0001) (Fig. 5A). Second, AE significantly reduced NfL serum levels in both sham and 6-OHDA groups (F_(1,29)_ = 32.6, *p*<0.0001), with 42% reduction in the sham group (NE = 232 ± 11, EX = 135 ± 11) and 25% reduction in the 6-OHDA group (NE = 328 ± 20, 6-OHDA NE = 245 ± 14) (Fig. 5A). There was no difference in NfL levels between sham NE and 6-OHDA EX groups (Fig. 5A), indicating the AE regimen reduced lesion-related NfL accumulation.

**Figure 5.**
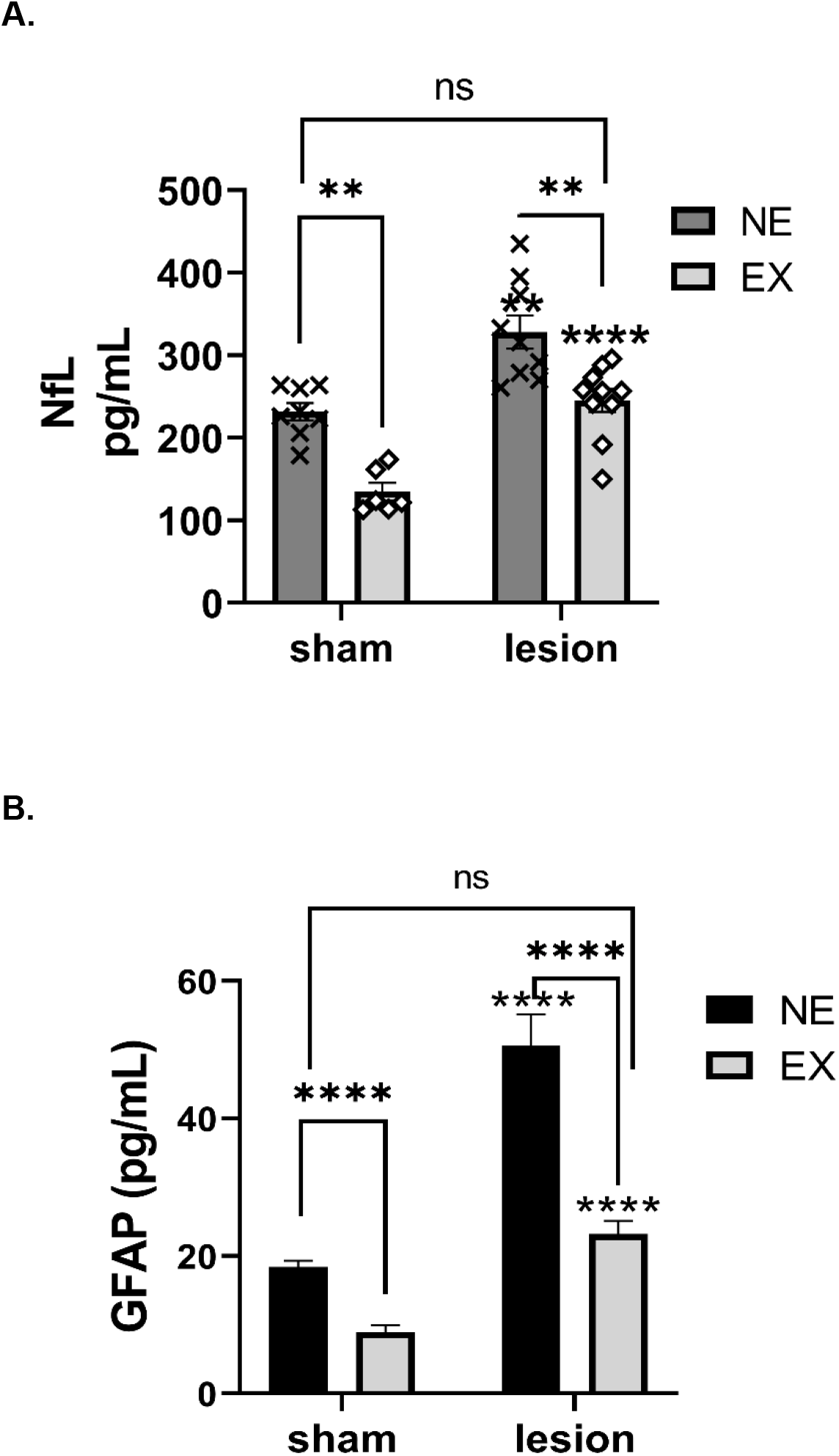
Serum NfL and GFAP levels. **A. NfL.** Lesion increased NfL levels and the AE regimen reduced NfL levels in the sham-operated and 6-OHDA groups. Unpaired t-test results; sham NE v lesion NE (t=3.20, \*\**p*=0.006, df=16), sham EX v lesion EX (t=8.00, ****p<0.0001, df=14), sham NE v sham EX (t=7.60, \*\*\*\**p*<0.0001, df=11), lesion NE v lesion EX (t=3.30, \*\**p*=0.004, df=19), sham NE v lesion EX (t=1.02, *p*=0.32, df=15).). **B. GFAP.** Lesion increased GFAP levels and the AE regimen reduced GFAP in the sham-operated and 6-OHDA groups. Unpaired t-test results; sham NE v lesion NE (t=5.60, \*\*\*\**p*<0.0001, df=13), sham EX v lesion EX (t=5.90, ****p<0.0001, df=15), sham NE v sham EX (t=7.0, \*\*\*\**p*<0.0001, df=11), lesion NE v lesion EX (t=5.70, \*\*\*\**p*<0.0001, df=17), sham NE v lesion EX (t=1.88, *p*=0.08, df=14).

GFAP serum levels also increased nearly 3-fold by 6-OHDA lesion in the NE group ((mean ± SEM) sham NE = 18.4 ± 0.90, 6-OHDA NE = 50.6 ± 4.6 (F_(1,28)_ = 60.4, *p*<0.0001)) (Fig. 5B). AE significantly reduced GFAP serum levels in both sham and 6-OHDA groups (F_(1,28)_ = 38.1, *p*<0.0001), with 52% reduction in the sham group (NE = 18.4 ± 0.90, EX = 8.9 ± 1.0) and 54% reduction in the 6-OHDA group (NE = 50.6 ± 4.6, 6-OHDA NE = 23.2 ± 1.9). There was no difference in GFAP levels between sham NE and 6-OHDA EX groups (Fig. 5B), indicating the AE regimen reduced lesion-related GFAP accumulation.

### Striatal dopamine content and tyrosine hydroxylase expression

There was an interaction of exercise x lesion (F_(1,33)_= 4.37, *p*<0.044), but no significant difference in DA loss ipsilateral to lesion between the NE (6.2 ng DA/mg protein, 98% loss) and EX (7.7 ng DA/mg protein, 97% loss) 6-OHDA lesioned groups (Fig. 6A). Sham-operation had a minimal effect on DA levels (Suppl Fig 7A).

**Figure 6.**
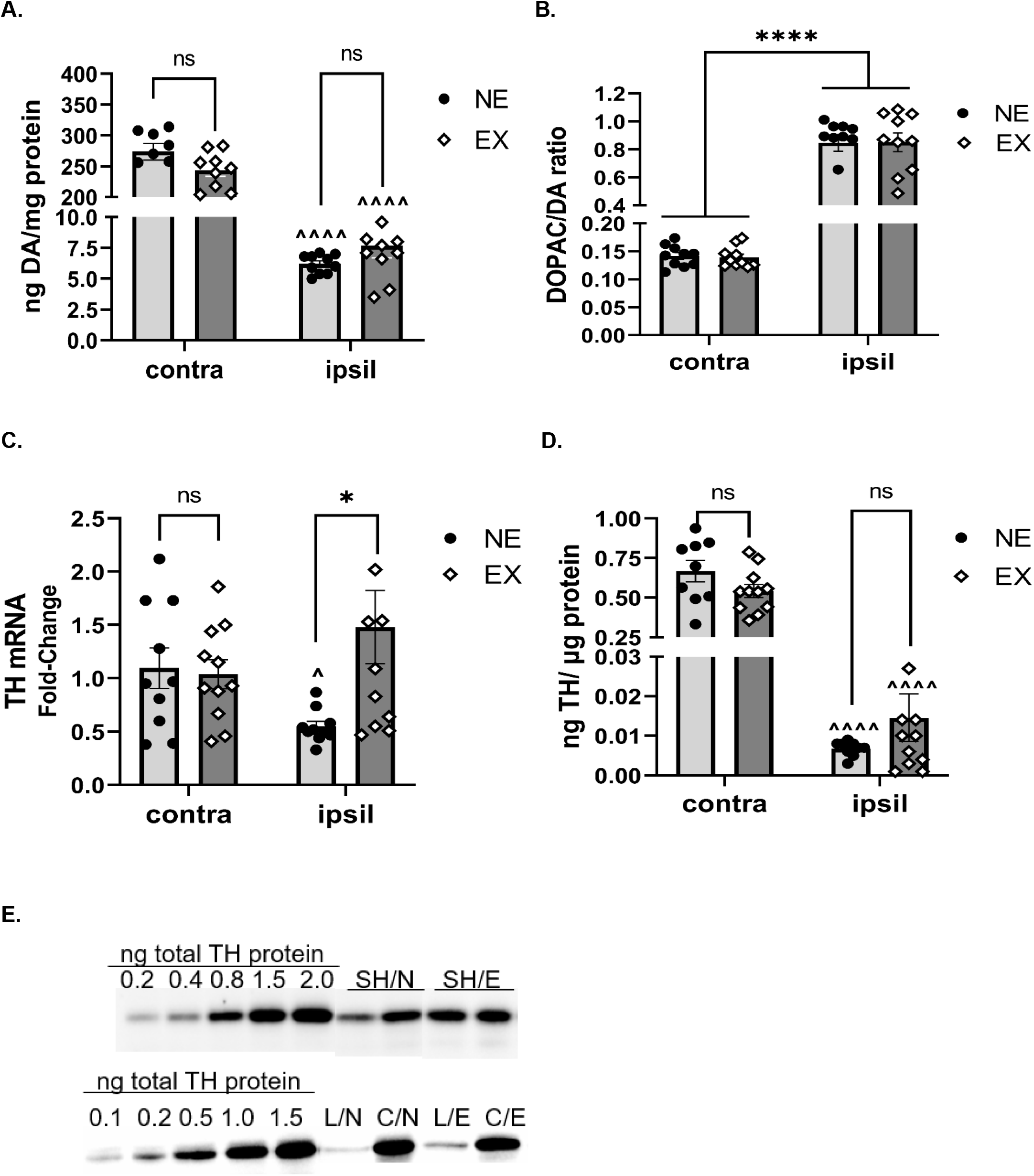
Exercise did not recover loss of striatal DA or TH protein. **A. DA loss.** As expected, lesion decreased DA tissue content more than 95% (F_(1,33)_= 814, *p <*0.0001). The AE regimen did not affect DA content respective to side of lesion (exercise, (F_(1,33)_= 3.59, *p*=0.067), exercise x lesion F_(1,33)_= 4.37, *p*=0.044). NE vs EX, contra (t=1.83, *p*=0.087, df=15); ipsil (t= 1.46, *p*=0.16, df=12). Contra vs ipsil, NE (t=19.6, ^^^^*p*<0.0001, df=7); EX (t=22.4, ^^^^*p*<0.0001, df=8). **B. DA turnover.** Lesion increased DA turnover ∼5-fold (F_(1,17)_= 256, *p <*0.0001). Exercise had no effect or interaction with lesion (exercise, (F_(1,19)_= 0.000, *p*=0.99), exercise x lesion F_(1,17)_= 0.01, *p*=0.93). **C. TH mRNA.** AE prevented lesion-related loss of mRNA (exercise x lesion, F_(1,19)_= 6.88, *p*=0.017). Lesion, F_(1,19)_= 0.078, *p* =0.78; exercise, F_(1,19)_= 3.33, *p* =0.08). Contra vs ipsil, NE (t=2.83, ^*p*=0.020, df=9); EX (t=1.41 *p*=0.19, df=10). NE vs EX, ipsil, (t=2.56, \**p*=0.019, df=19), contra (t=0.25, *p*=0.80, df=19); **D. TH protein expression.** Lesion decreased TH (ng TH / µg protein) expression >95% (lesion F_(1,17)_= 256, *p <*0.0001). The AE regimen did not affect TH protein between groups respective to side of lesion (exercise, (F_(1,19)_= 2.30, *p*=0.14), exercise x lesion F_(1,17)_= 3.26, *p*=0.088). Contra vs ipsil, NE (t=9.8, ^^^^*p*<0.0001, df=8); EX (t=13.6, ^^^^*p*<0.0001, df=10). **E. Representative quantitative western blot of total TH protein.** TH was quantified against a calibrated total TH standard, representing the ng quantities of TH protein loaded for the standard curve. Sham (SH) groups – non-exercise (N) and exercise (E) groups and 6-OHDA groups (L-Lesioned (Ipsilateral)/C – Contralateral hemispheres). Due to the significant reduction in TH expected in the lesioned hemispheres, 10 µg total protein was loaded from lesioned hemispheres and 2 µg loaded from contralateral hemisphere (C)).

The 6-OHDA lesion increased DA turnover ∼5-fold (F_(1, 17)_ = 256, *p*<0.0001) (Fig 6B). Exercise had no effect or interaction with lesion on increased striatal DA turnover due to lesion. Sham-operation had no effect on DA turnover (Suppl Fig 7B).

TH mRNA expression in striatum was reduced ∼50% by 6-OHDA in the NE group, and this difference was not observed in the EX group (lesion x exercise, F_(1,19)_= 6.88, *p*=0.016) (Fig. 6C). However, exercise alone had no effect on TH mRNA expression in the 6-OHDA group (exercise, F_(1,19)_= 3.33, *p*=0.084) or in the sham group (fold-change, mean ± SEM; NE=0.73 ± 0.15, EX=0.85 ± 0.12) (t=0.645, *p=*0.53).

As expected with the 6-OHDA hemi-lesion, there was major (>90%) loss of TH protein expression (lesion F_(1,18)_= 256, *p*<0.0001). Despite alleviation of motor impairment, TH protein expression was unaffected by AE ipsilateral or contralateral to lesion (exercise, F_(1,18)_= 2.30, *p*=0.15) (Fig. 6D). Notably, AE did increase TH protein expression in the sham-op group (exercise, F_(1,24)_= 11.4, *p*=0.003) (Suppl Fig 7C).

## Discussion

This proof-of-concept study indicates that an AE regimen, which begins after establishing motor impairment and emulates exercise intensity, duration and frequency within the physical capabilities of early-stage patients (Salvatore et al., 2022), promotes recovery of established motor impairment and mitigates severity of hypokinesia. These motor improvements were associated with decreased levels of serum NfLand GFAP, emerging as blood-based biomarkers of disease severity in PD (Buhmann et al., 2023; Youssef et al., 2023). With these translational elements of AE parameters and timing of intervention in place, these results represent a major step forward in that increased striatal TH protein expression or DA tissue levels may not be required for AE-related motor recovery. However, AE impact may be monitored peripherally by a serum biomarker. We point out that the timing of AE intervention also coincided with >80% loss of TH and DA in the striatum (Kasanga et al., 2022). This also improves translation potential to human PD, since ≥ 80% TH loss is present at the time of parkinsonian sign onset (Bernheimer et al., 1973, Bezard et al., 2001; Kordower et al., 2013). Therefore, this design enabled us to not only determine if the AE regimen could alleviate motor recovery, but also whether striatal TH or DA loss could recover to some extent contemporaneously in conjunction with alleviation of impairment. We did not directly evaluate DA release capacity following AE in striatum (currently under investigation). However, our current microdialysis results indicate the potential for DA release is practically abolished under depolarizing stimulation; being directly tied to DA tissue content. To this end, our results are expected to help resolve the ambiguity as to why AE can alleviate onset of motor impairment with (Choe et al., 2012; Lau et al., 2011; Real et al., 2017; Smith et al., 2011; Tajari et al., 2010; Yoon et al., 2007) or without (Arnold et al., 2016, 2017; Al-Jarrah et al., 2007; Chen et al., 2018; Fischer et al., 2004; Gorton et al., 2010; Hood et al., 2016; O’Dell et al., 2007; Petzinger et al., 2007; Tillerson et al., 2003; Tsai et al., 2019) any commensurate change in striatal DA or DA markers.

AE is being realized as a top intervention strategy for delaying motor symptom onset during the prodromal phase of PD (Daalen et al., 2022). Substantial evidence from rodent PD studies support that neuroprotection by AE is feasible when initiated prior to or within several days of nigrostriatal lesion (Table 1). In fact, the duration of exercise conducted pre-lesion may be correlative to the level of neuroprotection (Gerecke et al., 2010). These results have translational value to the human condition, as retrospective and prospective studies lend strong support that a physically-active lifestyle reduces the risk of PD (Portugal et al., 2023; Uhrbrand et al., 2015; Yang et al., 2015). However, since the prodromal phase PD still lacks a definitive signature of behavioral phenotype and biomarker profile to date, human studies of AE can only evaluate its impact on a trajectory from diagnosis forward. This brings up an important and critical point in interpreting and translating outcomes of preclinical back to human PD in that the vast majority of preclinical studies began AE either 1-4 weeks before lesion induction or within just a few days after nigrostriatal lesion induction (Table 1). Arguably, with no loss of striatal TH or DA pre-lesion or little loss within a few days after lesion in some rodent studies, the modeling of lesion severity in human PD at motor symptom onset by pre- or early AE intervention is not represented. Thus the translational potential of neurobiological mechanisms of AE in rodent models with pre- or very early post-lesion AE intervention would seem to be applicable to the human condition only in those with a physically-active lifestyle or in the prodromal phases of PD.

To our knowledge, this is the first study to date to evaluate potential motor recovery with AE intervention beginning after impairment has been established and is coincident with loss of striatal DA comparable to that in human PD, and incorporates translatable elements in duration, frequency, and intensity that are practicable in human PD. Roughly ∼40% of studies began exercise at least 5 days after lesion induction (Table 1), but far fewer of these established severe striatal DA marker loss by the lesion paradigm or motor phenotype before or early after exercise began (Chen et al, 2018; Petzinger et al, 2007; Poulton and Muir, 2005; Toy et al., 2014), and only one verified motor impairment (Wang et al., 2013). Petzinger and colleagues (2007) demonstrated increased motor function with exercise (5d/wk for 28 days), using rotarod-based assessment, without increased striatal DA tissue content or TH protein expression. However, motor impairment could not be established with this MPTP model, despite major loss of tissue DA. Nonetheless, this study verified significant dopamine depletion at 2 time points pre- and post-exercise, with no striatal DA recovery evident with exercise. Two studies did verify evidence of motor impairment either before or one wk after initiating exercise (5x/wk) and reported improvements in motor parameters in the EX group beginning the 2^nd^ week of exercise (Chen et al., 2018; Wang et al., 2013); notably a similar time point of AE effects seen in our study. Chen and colleagues reported greater DA release in EX lesioned rats, but notably, the release was still substantially lower (20% of release in non-lesioned rats), suggesting the increase in striatal DA release was likely not driving motor improvements. Choe and colleagues (2012) showed recovery of TH loss in striatum, but notably TH loss in the NE group was ∼50%, not near the severity of TH loss shown in association with the forelimb deficits in our study (Kasanga et al., 2022) or in patients at onset of motor impairment, nor did they report motor function. Our study builds upon the timing of AE impact beginning ∼ 2 wks after AE (Chen et al., 2018; Wang et al., 2013), but also confirms motor impairment established prior to exercise can be recovered using translatable elements within the capabilities of a PD patient (Kasanga et al., 2021; Salvatore et al., 2022) and does not obligate recovery of DA or TH loss comparable to a magnitude similar to that of a newly diagnosed PD patient.

A landmark study of post-mortem tissue by Kordower and colleagues (2013) clearly shows that there is at least 70-90% loss of striatal DA markers within 3 yrs of PD diagnosis, conifrming the conclusions of earlier work (Berheimer et al., 1973), and quantitative metrics in primate PD studies that 80% loss in striatum was associated with onset of motor impairment (Bezard et al., 2001). Moreover, little if any DA markers remain 4 yrs post-diagnosis. Our study suggests such profound loss in striatum is not recoverable by AE, but other studies indicate loss may be preventable if AE is initiated sometime before motor impairment. Even so, the results indicate recovery of lost DA markers in striatum is not required for motor improvement. This finding is in alignment with findings from other human and preclinical studies, outside the scope of exercise evaluation, that show recovery nor presence of normal levels of DA markers in striatum predict parkinsonian sign rescue or prevention (Blesa et al., 2012; Dave et al., 2014; Gash et al., 1996; Kordower et al., 2017; Perez-Taboada et al., 2020; Salvatore et al., 2016, 2017; 2019; 2023). Our results should help dissolve the perseverating focus that restoring DA loss in the striatum is necessary for motor improvements or preventing motor decline.

### Study strengths and limitations

There are 4 limitations in our study. Our study used young rats. As aging is the number one risk factor for PD, inclusion of aged rats would bring the relevant neurobiological background in most patients. Accordingly, we reported advanced aging, in the absence of nigrostriatal lesion, could diminish AE efficacy (Arnold et al., 2017), but on the other hand, AE was effective in rats that were 18 months old, which in humans roughly corresponds to 50-60 yr old (Quinn, 2005).

Therefore, this study sets the stage to examine efficacy of this AE regimen in older rats. The length of time of AE evaluation in our study was for 3 wks, corresponding to ∼20 human months (Quinn, 2005). Whereas motor benefit phenotypes were captured in this time frame, it will be necessary to determine the duration of AE impact. In fact, a training period of at least 6 months in human PD reduces UPDRS-III scores (Mak et al., 2017). In the same vein, it may be possible that striatal DA or TH levels be partially recovered with longer application of AE. Third, our study did not evaluate gait parameters, which are improved by AE (Chen et al., 2018; Hood et al., 2016; Tsai et al., 2019). It will be important to evaluate impact on gait using the same paradigm. In rodents, treadmill exercise causes fatigue (Wang and Wang, 2018; Zaretsky et al., 2018). In PD patients, fatigue occurs in 36% of PD patients (Zhou et al., 2023), can specifically reduce distance travelled in motor tests (Carvalho et al., 2020), and can be elicited by physical exertion in 20% of patients (Lin et al., 2021). Given the evidence of AE-related fatigue immediately following the first week of AE in both sham and 6-OHDA groups, an AE benefit on hypokinesia prevention may have been masked. The minor, but not significant, increase in forelimb use 1 wk after AE intervention suggests this possibility. Hypokinesia onset in this model occurs 21 days after lesion (Kasanga et al., 2022) and this timing was replicated in the NE group as compared to the NE group in the sham cohort. This onset corresponds to the end of the 2^nd^ wk of AE timing wise, and thus relative to locomotor activity vs the 1^st^ wk after AE, hypokinesia onset was prevented, as only the NE group in the lesioned cohort continued to worsen.

In addition to the 3 components of translation in the AE regimen, there are additional strengths of this study. Other than confirming exercise ability in all rats prior to random assignment to experimental groups, the rats were essentially sedentary. A substantial proportion of PD patients can be classified having a sedentary lifestyle (von Rosen et al., 2021), and may spend up to 98% of their day on sedentary to light intensity activity levels (Dontje et al., 2013). Therefore, if barriers to exercise can be overcome in sedentary patients (Afshari et al., 2016; Ellis et al, 2013; Schootemeijer et al., 2020) our results suggest that these individuals may still realize AE benefits. Another strength for translation potential is the simplicity and predictability of the AE protocol, which can be recreated quickly with relative ease, minimal equipment, and low risk for human PD treatment interventions. With the cost effectiveness and wide availability of AE equipment and health monitoring devices, this puts control into the hands of the patient, family, caregiver, and healthcare provider to give specific advice (even via telehealth) that a patient can safely take immediate action. This is an important factor when considering the possibility that neuroprotection by AE might be feasible when initiated in a defined prodromal period, which is still being elucidated (Nejtek et al., 2021; Portugal et al., 2023;Tolosa et al., 2021). Obtaining a referral to an outpatient therapy clinic or PD specific training program can take anywhere from 1 week to 1-2 months. Therefore, best practice guidelines for early onset PD in the future may shift to include encouragement of the appropriate patient to begin specific and trackable AE protocols even prior to starting a defined therapy or training program. Finally, our results report for the first time evidence that exercise efficacy may be obtained from a blood-based biomarker. Nigrostriatal lesion augmented Nfl and GFAP levels in serum, consistent with motor impairments and recent human literature that both biomarkers may correlate with disease severity (PD (Buhmann et al., 2023; Mollenhauer et al., 2020; Tang et al., 2023; Ye et al., 2021; Ygland Rodstrom et al., 2022; Youssef et al., 2023). The levels of both biomarkers decreased significantly in rats in the exercise group. Both biomarkers also decreased in the sham-group, which illuminates a new avenue to gauge exercise impact in the CNS by a peripheral biomarker.

### Summary

Previous rodent model studies strongly support that AE is neuroprotective, consistent with observations that physically active humans incur less risk of PD. As AE is garnering priority as a strategy to alleviate motor impairment with disease progression (Alberts et al., 2016; Daalen et al., 2022; Mak et al., 2017; Ridgel and Ault, 2019; Schenkman et al., 2018), it is imperative to apply exercise parameters consistent with PD patients’ physical capabilities and evaluate efficacy after motor impairment is established. Our study has demonstrated a critical proof-of-concept in this regard in that a moderate intensity regimen applied 3 times weekly promotes recovery and alleviates onset of 2 specific motor impairments; forelimb use recovery and alleviation of hypokinesia severity. These AE-driven increases in motor function were accompanied by reduced serum levels of NfL and GFAP, both recently shown to be peripheral blood-based biomarkers of disease severity in human PD. Despite these indications of AE efficacy, striatal TH and DA loss were not recovered, indicating AE mechanisms may not require a change in DA biosynthesis or release, as severe loss of tissue DA essentially eliminated DA release capacity at the time AE was initiated. This study serves as a platform upon which to interrogate neurobiological mechanisms driving AE impact on motor function, verify AE efficacy with blood-based biomarkers, and, accordingly, increase translation to gauge exercise efficacy in human PD.

## Supporting information

Kasanga et al bioRxiv July 2023

## Acknowledgements

**Funding:** This work was supported by the Department of Defense Parkinson’s Research Program, Investigator-Initiated Research Award (W81XWH-19-1-0757) award to MFS and R01ES033892 and RF1NS130713 to JRR. EAK was supported by the Office of Vice President for Research and Innovation, the Institute for Healthy Aging, and National Institutes of Health/National Institute on Aging Predoctoral International Fellowship (T32 AG020494), and the Parkinson’s Foundation Visiting Scholar Award. The funders had no role in the study design, collection, analysis, and interpretation of data, writing of the manuscript, or decision to submit the article for publication.

