## Supplementary material for "Moderate intensity aerobic exercise in 6-OHDA-lesioned rats alleviates established motor deficits and reduces neurofilament light and glial fibrillary acidic protein serum levels without increased striatal dopamine or tyrosine hydroxylase protein": Kasanga et al bioRxiv July 2023

**Supplemental Data**

**Suppl Table 1**

### Study criterion for statistical analysis:

1) **6-OHDA lesion efficacy verified by >25% loss of forelimb use. AMPH rotation is secondary validation of lesion, both 7 days post lesion.**
2) **Rats continue in assigned group, regardless of FAS outcome**
3) **Compare striatal DA and TH loss after study vs FAS results from day 7.**
4) **Grubb's test on negative day 7 FAS results (<25% loss), if >90% DA loss. Include result if not outlier.**
5) **All DA, TH protein, and NfL and GFAP results represent confirmed >75% loss of DA and TH.**

**Suppl Table 2. Correlation of body weight and pre- and post-locomotor assessments**

| | Pre-Exercise Assessment | | Post-Exercise Assessment | | | |
|---|---|---|---|---|---|---|
| Parameters | Pearson's Coefficient | *p*-value | Pearson's Coefficient | *p*-value | Pearson's Coefficient | *p*-value |
| FAS | -0.2298 | 0.3590 | 0.0425 | 0.9204 | 0.2020 | 0.6022 |
| Total Distance | 0.0292 | 0.9056 | -0.0904 | 0.8171 | 0.4209 | 0.2593 |
| Speed | -0.1257 | 0.6082 | -0.3798 | 0.3133 | 0.1862 | 0.6316 |

**Suppl Fig. 1**

**A.**

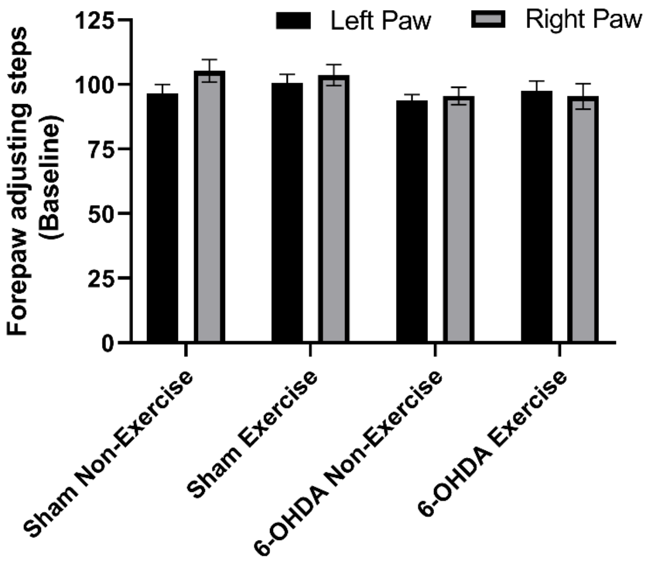

**B.**

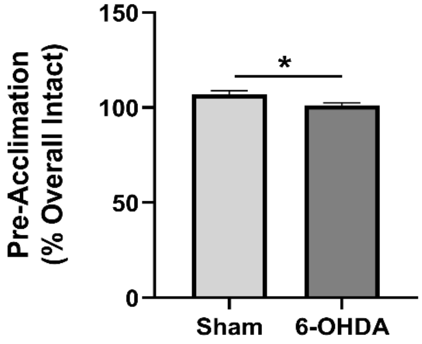

**C.**

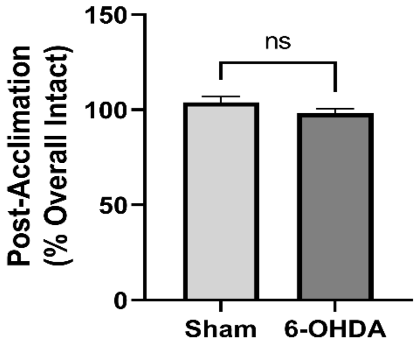

**Supplemental Figure 1. Forelimb use prior to group assignments and before and after exercise acclimation. A. Overall forelimb use.** Prior to eventual group assignment and exercise acclimation, there were no differences in right or left forelimb use. Left v right ($F_{(1, 68)} = 1.02$, $p$ =0.32) or group assignment ($F_{(3, 68)} = 1.54$, $p$ =0.21). **B. Forelimb use (Right/Left), prior to exercise acclimation.** There was ~5% difference in percentage of right forelimb use in rats eventually assigned to sham (106.9 ± 2.1) % or 6-OHDA (101.2 ± 1.4%) groups (t=2.41, *$p$ =0.021, df=36). **C. Forelimb use (Right/Left), after exercise acclimation.** There was no significant difference in rats eventually assigned to sham (103.9 ± 3.0%) or 6-OHDA (98.3 ± 2.2%) groups (t=1.54, $p$ =0.131, df=36).

**Suppl Fig. 2**

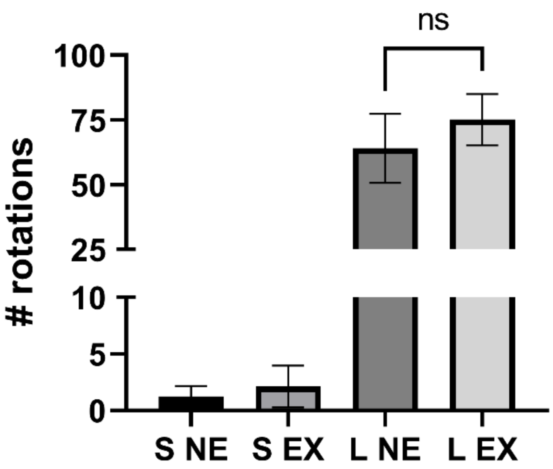

**Supplemental Figure 2. AMPH-rotations day 7 post-lesion.** As expected, there was a highly significant difference in rotation number between sham-operated (S) and 6-OHDA lesioned (L) groups ($F_{(3,35)}$ =15.8, $p$<0.0001). There was no significant difference in AMPH rotation number between rats assigned into the NE or EX groups 7 days after 6-OHDA (t=0.87, $p$=0.392).

**Suppl Fig. 3**

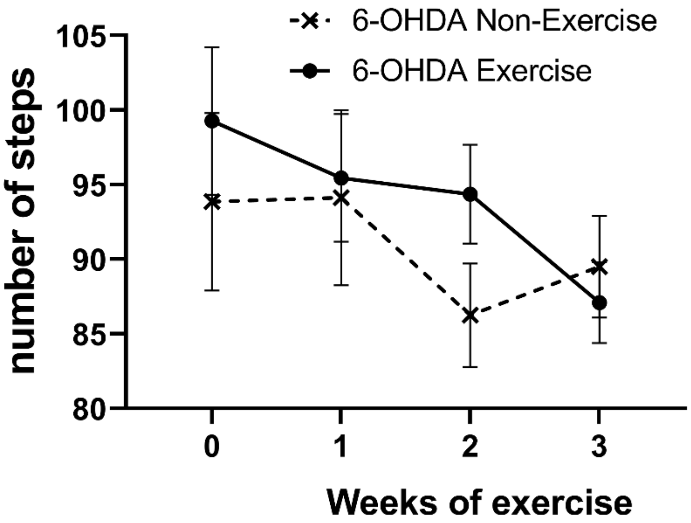

**Supplemental Figure 3. Exercise had no impact on forelimb use associated with the intact side.** Exercise ($F_{(1,17)}$ =0.27, $p$=0.60). Duration of exercise x exercise ($F_{(3,50)}$ = 1.94, $p$ = 0.13). weeks post-lesion ($F_{(3,50)}$ = 6.09, $p$ = 0.001).

**Suppl Fig. 4**

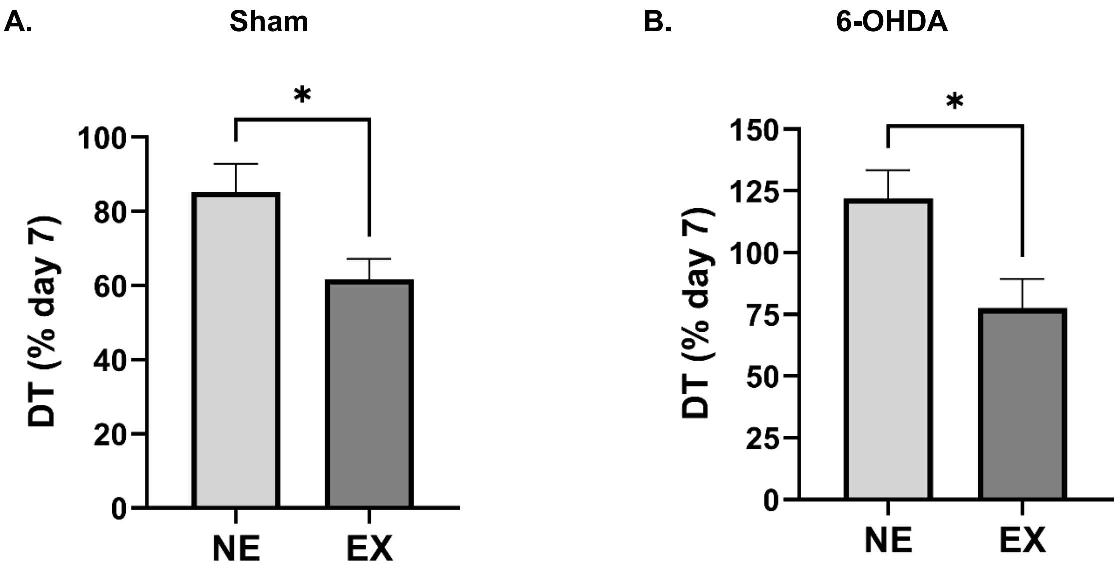

**Supplemental Figure 4. Evidence of fatigue with exercise.** As compared to locomotor activity in the NE groups, the EX group in the **A)** sham-operation (t=2.52, *p* =0.027, df=12) and **B)** 6-OHDA groups (t=2.70, *p* =0.014, df=19) exhibited a significant decrease in locomotor activity the day after completing the first week of exercise.

**Suppl Fig. 5**

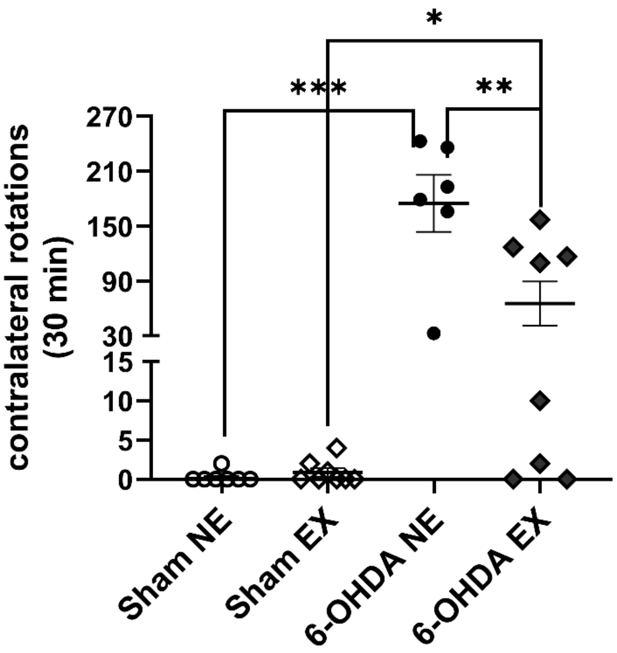

**Supplemental Figure 5. Apomorphine rotations.** There was a highly significant difference in the number of rotations contralateral to lesion among the 4 groups ($F_{(4,30)}$ = 19.6, *p*=0.0002). Mann-Whitney test comparisons; Sham NE v 6-OHDA NE (Sum of ranks, 36, 69, ****p*=0.0003 U=0), Sham EX v 6-OHDA EX (Sum of ranks, 48.5, 87.5, **p*=0.034, U=12.5), 6-OHDA NE v 6-OHDA EX (Sum of ranks, 65, 40, ***p*=0.008, U=4).

**Suppl Fig. 6**

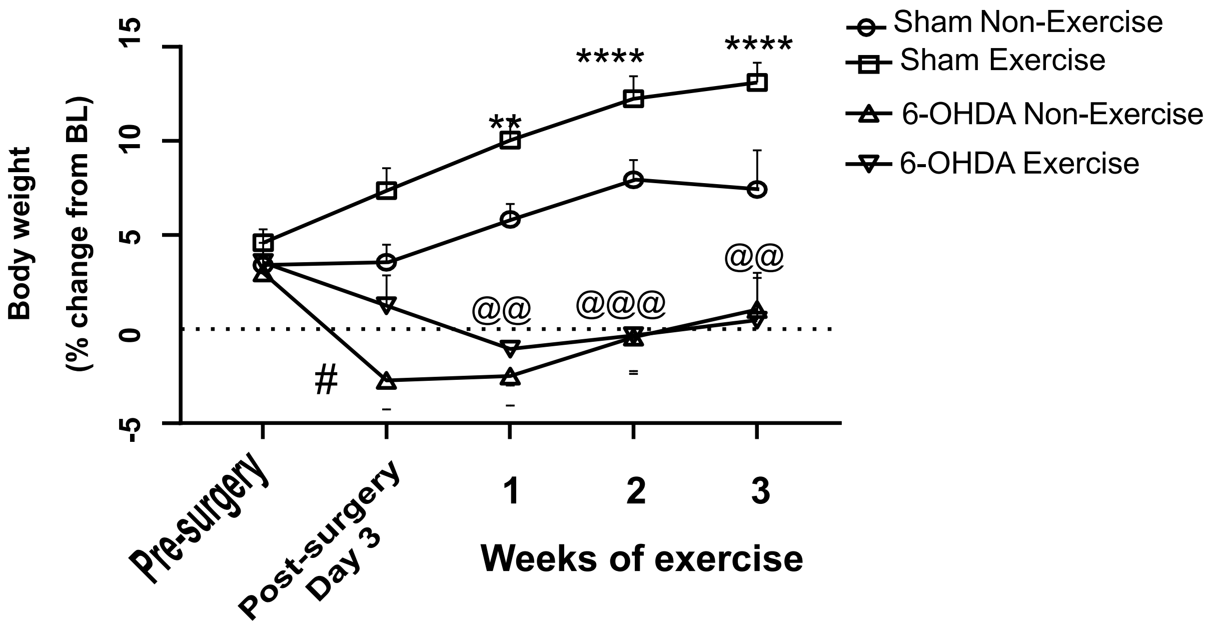

**Supplemental Figure 6. Impact of exercise on body weight.** There was a significant difference between groups ($F_{(3,\ 31)}$=12.95, $p$<0.0001), time post-exercise ($F_{(2.5,\ 75.6)}$=7.84, $p$<0.001) and group x time post-exercise interaction ($F_{(12,122)}$=7.56, $p$<0.0001). Animals receiving 6-OHDA showed a significant decline in body weight 3 days after surgery (# $p$<0.05 - Post-surgery day 3 – Sham vs 6-OHDA groups). This decline in body weight was sustained throughout the study (@ all $p$ ≤0.01,- Sham Exercise vs 6-OHDA Exercise). An increase in body weight was also observed in the exercise group starting from exercise post-session 1 to 3 (* all $p$≤0.01 - Post-surgery vs Exercise sessions (Sham Exercise)).

**Suppl Fig. 7**

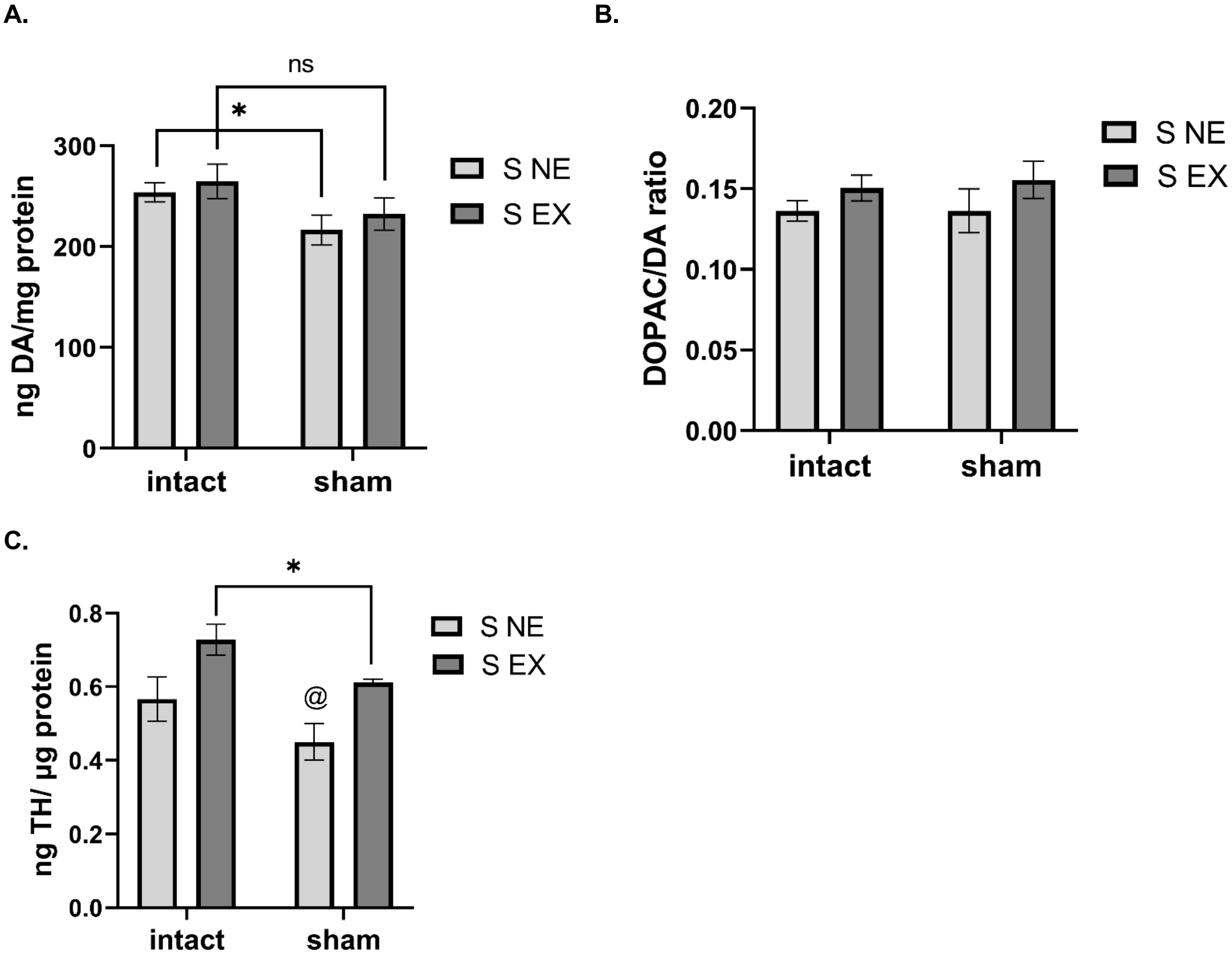

**Supplemental Figure 7. Sham-operation and striatal DA tissue content, DA turnover and TH protein expression. A. DA tissue content.** Sham-operation ($F_{(1,24)}$= 5.80, *p*=0.024), exercise ($F_{(1,24)}$= 0.86, *p*=0.36), sham-op x exercise ($F_{(1,24)}$= 0.031, *p*=0.86). Intact v sham; NE (t=2.21, **p*=0.048, df=12), EX (t=1.38, *p*=0.19 , df=12). **B. DA turnover.** Sham-operation ($F_{(1,12)}$= 0.05, *p*=0.82), exercise ($F_{(1,14)}$= 2.62, *p*=0.13), sham-op x exercise ($F_{(1,12)}$= 0.11, *p*=0.75). **C. TH protein.** Sham-operation ($F_{(1,24)}$= 5.85, *p*=0.024), exercise ($F_{(1,24)}$= 11.4, *p*=0.003), sham-op x exercise ($F_{(1,24)}$= 0.00, *p*=0.99). Intact v sham; NE (t=1.45, *p*=0.17, df=13), EX (t=2.53, **p*=0.028, df=11). NE v EX; intact (t=2.14, *p*=0.052, df=13), sham (t=2.96, $^{@}$*p*=0.013 , df=11).

### Additional Method Details

*Microdialysis*

On day of microdialysis, probes for the striatal site extended 3 mm beyond the cannula tip (molecular weight cut-off 100 kDA) and the nigral probes (results to be used for another study) extended 1 mm beyond (molecular weight cut-off 20 kDA). In both sites, artificial cerebral spinal fluid (aCSF; 147 mM NaCl, 2.8 mM KCl, 1.2 mM CaCl2, 1.2 mM MgCl2, pH 7.40 ± 0.02, sterile filtered) were perfused with a syringe pump (model 400, CMA Microdialysis) at 2 µL/min.

*UHPLC analysis*

UHPLC-ECD mobile phase MD-TM consisted of 75 mM sodium dihyrdogen phosphate monohydrate, 1.7 mM 1-Octanesulfonic Acid sodium salt, 100 µL/L Triethylamine, 25 µM EDTA, 10% Acetonitrile, at a pH of 3.00 (Fisher Scientifi)c. The UHPLC pump (Ultimate 3000BM) was run with a 0.6 mL/min isocratic gradient at ~255 Bar and coupled to an Ultimate 3000RS electrochemical detector and set to DC mode. Two cells were used, a 6020RS omni coulometric cell set at +300 mV to oxidize any contaminants in the mobile phase, and a 2-channel 6011RS ultra analytical cell set at -150 mV (Channel 1) and +220 mV (Channel 2). The analytical column that was used was a BDS Hypersil (150mm X 3mm) C18 column with a particle size of 3µm operating at 37° C. Samples and standards were placed in a WPS 3000TBRS autosampler maintained at 4 ˚C during analysis. Injection volume for all standards and samples was 10 µL. The injection needle was rinsed with 10% degassed MeOH between injections. For analysis of DA tissue content, a standard curve ranging from 1.56 ng/mL to 1200 ng/mL (1.56, 3.13, 6.25, 12.5, 25, 50, 100, 200, 400, 800, 1200) was used and for analysis of microdialysis samples, the standard curve ranged from 0.05-75 ng/ml (0.05, 0.1, 0.25, 0.50, 0.75, 1.0, 1.5, 2.5, 5, 7.5, 10, 15, 25, 50, 75) and correlation coefficients met or exceeded 0.99 and values obtained from all samples fell within the range of the standard curves.

*mRNA extraction and analysis*

iScript™ cDNA Synthesis Kit mixture consisted of 1 µL of RNA template, 4 µL of the 5x reverse-transcription reaction mix, 14 µL of nuclease-free water, and 1 µL of iScript™ reverse transcriptase for a total reaction volume of 20 µL. PCR cycling conditions consisted of four steps; 1) the priming step was set at 25°C for 5 minutes, 2) the reverse transcription step at 46oC for 20 minutes, 3) the reverse transcriptase inactivation step at 95oC for 1 minute, and 4) a hold at 4° C. The reaction mixture for TH mRNA quantitation had a total volume of 20 µL and consisted of 10 µL of SsoAdvanced Universal SYBR® Green Supermix, 1µL of PrimePCR™ SYBR® Green Assay primer mixture, varying amounts of nuclease-free water, and varying amounts of cDNA template in the wells of a 96-well plate. The primer mixes used for this experiment were the PrimePCR™ SYBR® Green Assay: TH, Rat for the gene of interest and the PrimePCR™ SYBR® Green Assay: GAPDH Rat for the control gene.

PCR cycling program was set for SYBR only and consisted of four steps; 1) the polymerase activation and initial DNA denaturation step at 95°C for 30 seconds, 2) the subsequent denaturation step at 95°C for 5 seconds, 3) the annealing/extension and plate read step at 60°C for 30 seconds, and 4) a melt curve ranging from 65°C to 95°C rising in increments of 0.5°C per step for 5 seconds each. The subsequent denaturation and annealing/extension and plate read steps were repeated for 40 cycles.

| (notes) | Year | PMID | Author | Model | Timing of aerobic Exercise Intervention Relative to Lesion | Frequency and Duration of Exercise | Exercise Intensity | Primary Motor Effect | Primary Neurochem Finding | |
|---|---|---|---|---|---|---|---|---|---|---|
| | 2003 | 12809709 | Tillerson | MPTP & 6-OHDA (rat and mice) | within 1st day of lesion | 2x/d for 9 d | 15 m/min, 15min | forelimb akinesia and placement asymmetry reduced | no changes in striatal TH or DA | |
| * at least 5 days after lesion | 2005 | 15817277 | Poulton | 10 μg 6-OHDA in MFB (F rat) | 1 wk post-lesion | 2x/d 6x/wk for 1 mo. | 13m/min, 40min | no effect on motor | preserved DA and DOPAC loss in str | |
| | 2007 | 17157992 | O'Dell | 3 μg 6-OHDA in str (rat) | 2.5 wks pre-lesion | forced 2x/day + voluntary in cage for 6.5 wks | max 10.5m/min | accelerated recovery of forelimb use | no changes in striatal TH+ cells or DAT | |
| | 2007 | 17644250 | Yoon | 20 μg 6-OHDA in str (rat) | 24 hr post-lesion | 14 d (consecutive) | 2-3m/min, 30min | decrease in apomorhpine rotations | enhanced survival of DA and preserved TH+ cells in SN | |
| * at least 5 days after lesion | 2007 | 17507552 | Petzinger | MPTP (mice) | 5 d post-MPTP regimen | 5x/wk for 28 d | 7.6-20.5m/min, 60min | improved motor coordiantion and balance | no effect on DA in SN or str | |
| | 2010 | 19900418 | Tajiri | 20 μg 6-OHDA in str (rat) | 24 hr post-lesion | 5x/wk for 4 wks | 11m/min, 30min | decrease in apomorhpine rotations and improved forelimb asymmetry | preserved TH fibers in str and TH+ cells in SN, increased BDNF and GDNF in str | |
| | 2010 | 20116369 | Gerecke | MPTP (mice) | 1, 2, and 3 mo. pre-lesion | 1, 2, or 3 mo. | voluntary running | n/a | no effect on DA in str, preserved DA loss in SN from 3mo of exercise | |
| * at least 5 days after lesion | 2010 | 20472000 | Gorton | MPTP (mice) | 5 d post-MPTP regimen | 5x/wk for 6 wks | 6.7-8.5m/min 30-60min | improved motor coordination and balance | no effect on DA levels in any brain region | |
| | 2011 | 21375602 | Lau | MPTP (mice) | 1 wk pre-lesion | 5x/wk for 18 wks | 15m/min, 40min | improved coordination and balance | preserved TH DA, and DAT in str and TH in SN, increase of BDNF and GDNF in SN and str | |
| * at least 5 days after lesion | 2012 | 23129977 | Choe | 20 μg 6-OHDA in str (rat) | 5 d post-lesion | 30m/d for 16 d | 10m/min | n/a | preserved TH neurons, increased TH expn in striatum | |
| | 2012 | 22231471 | Dutra | 10 μg 6-OHDA in MFB (rat) | 3 d post-lesion | 5d/wk for 4 wks | (N/A)m/min, 20 up to 60min | forelimb akinesia reduced and improved balance | no effect on TH neurons, decrease of GFAP in str | |
| * at least 5 days after lesion | 2013 | 24278884 | Cho | 20 μg 6-OHDA in str (F rat) | 4 wks post-lesion | 1x/d for 14 d | 8m/min, 30min | n/a | preserved TH expn in SN and str | |
| * at least 5 days after lesion | 2013 | 24278239 | Wang | 40 μg 6-OHDA in b/l str (rat) | 2 wks post-lesion | 5d (consecutive)/wk for 4 wks | ~5-6m/min, 20min | imrpoved motor coordination, balance, rearing counts and spontaneous activity | greater loss of TH in SN and str from exericse | confirmed motor impairment before exercise |
| | 2014 | 25264157 | Tuon | 8 μg 6-OHDA in str (mice) | 8 wk pre-lesion | 3-4d/wk for 8 wks | 13-17m/min, 50min | no effect on distance traveled yet decrease in apomorphine rotations | increased proBDNF in str and hippocampus | |
| * at least 5 days after lesion | 2014 | 24090962 | Goes | 5 μg 6-OHDA in str (mice) | 1 wk post-lesion | 5x/wk for 4 wks | swimming | improved motor coordination and balance | preserved DA, HVA and DOPAC loss in str | |
| * at least 5 days after lesion | 2014 | 24239657 | Landers | 10 μg 6-OHDA in MFB (rat) | 1 wk post-lesion | 5x/wk for 4 wks | 15m/min, 30min | no effect on motor | n/a | |
| * at least 5 days after lesion | 2014 | 24316165 | Toy | MPTP (mice) | 5 d post-MPTP regimen | 5x/wk for 6 wks | 10-24m/min | n/a | no effect on DA in str | |
| * at least 5 days after lesion | 2015 | 25943481 | Sconce | MPTP (mice) | 2 wks post-MPTP regimen | voluntary running in cage | voluntary running | improved grip strength and gait pattern | no effect on TH in SN or str | |
| | 2016 | 27350080 | Hood | MPTP | 1 wk pre-MPTP regimen | 5x/wk for 4 wks | 11m/min, 60min | increase in spontaneous locomotion w/ or w/o MPTP | no effect on striatal TH or DAT | |
| | 2017 | 28713483 | Da Costa | 12 μg 6-OHDA in str | 24 hr post-lesion | 14 d (consecutive) | 12m/min, 30min | improved motor coordianation and decreased apomorphine rotations | decreased DA loss in str, increased BDNF in str, PFC, and hippocampus | |
| | 2017 | 28801819 | Real | 6 μg 6-OHDA in str | 30 d pre-lesion | 3d/wk for 30 d | 10m/min, 40min | *decrease in apomorphine rotations | decrease of GFAP in SN and str, preserved TH in SN and str, | |
| | 2017 | 29204298 | Chen | 4 μg 6-OHDA in str | 24 hr post-lesion | 4 wks | 11m/min, 30min | forelimb placement asymmetry reduced | reduced striatal neuron spantaneous firing | |
| * at least 5 days after lesion | 2017 | 28243821 | Garcia | 12 μg 6-OHDA in str (rat) | 1, 2, 3, 4 mo. post-lesion | 3x/wk up to 4 mo. | 10m/min, 40min | n/a | preserved TH loss in SN and str after 3-4 mo. of exercise | |
| * at least 5 days after lesion | 2017 | 28754312 | Churchill | MPTP (mice) | 4 wks post-MPTP regimen | 5x/wk for 4 wks | 10.8m/min, 60min | improved motor coordination and gait performance | no effect on TH in SN or str | |
| * at least 5 days after lesion | 2018 | 29507426 | Chen | 6-OHDA (rat) | 1 wk post-lesion | 5d (consectuive)/wk for 4 wks | 11m/min, 30min | improved coordination and balance, walking speed, and hindlimb support and decrease in apomorphine rotations | enhanced DA release and acute recovery of cortico-striatal plasticity | |
| * at least 5 days after lesion | 2018 | 29019056 | Klemann | MPTP (mice) | 3 wks post-MPTP regimen | 2x/d for 28 d | 12m/min, 60min | improved motor corrdination and balance | no effect on TH+ cells in SN or VTA | |
| | 2019 | 29271291 | Real | 6 μg 6-OHDA in str (rat) | 2 d post-lesion | 3x/wk | 10m/min, 40min | no effect on distance traveled yet forelimb placement asymmetry reduced at day 9 but lost at day 29 | no effect on GFAP in SN with decrease of GFAP in str at day 30, preserved TH+ cells in SN and str at day 30 | |
| | 2019 | 31885725 | Tsai | 6 μg 6-OHDA in MFB (rat) | 2 wks pre-lesion and immediately post-lesion | 8 wks | voluntary running | *decrease in apomorhpine rotations and improved gait pattern | preserved TH+ cells in SN, no effect on str TH | |
| *6-OHDA exercise + blueberry juice (no 6-ohda + exercise only group) | 2022 | 353212527 | Castro | 6 μg 6-OHDA | 4 wks pre-lesion | 8 wks | voluntary running | decrease in apomorhpine rotations | no effect on TH in str, preserved TH+ cells in SN, no effect on GDNF in SN but exercise alone increased GDNF in str | |
